# “Enteric glia as a source of neural progenitors in adult zebrafish”

**DOI:** 10.1101/2020.02.14.949859

**Authors:** Sarah McCallum, Yuuki Obata, Evangelia Fourli, Stefan Boeing, Christopher J Peddie, Qiling Xu, Stuart Horswell, Robert Kelsh, Lucy Collinson, David Wilkinson, Carmen Pin, Vassilis Pachnis, Tiffany Heanue

## Abstract

The presence and identity of neural progenitors in the enteric nervous system (ENS) of vertebrates is a matter of intense debate. Here we demonstrate that the non-neuronal ENS cell compartment of teleosts shares molecular and morphological characteristics with mammalian enteric glia but cannot be identified by the expression of canonical glia markers. However, unlike their mammalian counterparts, which are generally quiescent and do not undergo neuronal differentiation during homeostasis, we show that a relatively high proportion of zebrafish enteric glia proliferate under physiological conditions giving rise to progeny that differentiate into enteric neurons. We also provide evidence that, similar to brain neural stem cells, the activation and neuronal differentiation of enteric glia are regulated by Notch signalling. Our experiments reveal remarkable similarities between enteric glia and brain neural stem cells in teleosts and open new possibilities for use of mammalian enteric glia as a potential source of neurons to restore the activity of intestinal neural circuits compromised by injury or disease.

## Introduction

Tissue integrity and repair depend on the regulated dynamics of adult stem cells, which share the capacity to replenish cellular compartments depleted by physiological turnover or disease. Studies on neural stem cells (NSCs) have advanced fundamental brain research and opened new and exciting opportunities for regenerative neuroscience (Morales and Mira, 2019). However, as NSC research has focused primarily on the central nervous system (CNS), our understanding of the homeostasis and regenerative potential of peripheral neural networks, and particularly the enteric nervous system (ENS), is minimal and at best phenomenological. This gap in knowledge impedes progress in fundamental gastrointestinal biology and stymies the generation of potential therapeutic strategies for repairing intestinal neural circuits with developmental deficits or damaged by injury or disease.

The ENS encompasses the intrinsic neuroglia networks of the gastrointestinal (GI) tract that are essential for digestive function and gut homeostasis (Furness, 2006). In vertebrates, assembly of the ENS begins during embryogenesis with invasion of the foregut by a small founder population of neural crest (NC) cells that proliferate and colonise the entire GI tract generating diverse types of enteric neurons and glial cells organised into networks of interconnected ganglia (Heanue and Pachnis, 2007). ENS development depends on the integrated activity of NC cell lineage-intrinsic programmes and signals from surrounding non-neuroectodermal gut tissues, which ultimately determine the organisation and physiological properties of intestinal neuroglial networks (Avetisyan et al., 2015; Rao and Gershon, 2018). Despite considerable progress in understanding the developmental mechanisms underpinning the assembly of intestinal neural circuits, much less is known about the dynamics of ENS cell lineages in adult animals, during homeostasis or in response to gut pathology. The predominant view holds that the vast majority of enteric neurons in the mammalian ENS are born during embryogenesis and early postnatal stages and remain functionally integrated into the intestinal circuitry throughout life (Bergner et al., 2014; Joseph et al., 2011; Laranjeira et al., 2011; Pham et al., 1991). Likewise, enteric glial cells (EGCs) are generally quiescent, with only a small fraction proliferating at any given time (Joseph et al., 2011; Kabouridis et al., 2015). Despite this static view of the ENS at homeostasis, lineage tracing experiments in mice have provided evidence that under experimental conditions, such as chemical injury of the ganglionic plexus and bacterial infection, a small fraction of Sox10^+^ and Sox2^+^ EGCs can differentiate into neurons (Belkind-Gerson et al., 2017; Belkind-Gerson et al., 2015; Laranjeira et al., 2011). However, a recent study has argued that a population of Sox10^-^Nestin^+^ ENS cells undergo extensive proliferation and neuronal differentiation even under physiological conditions, replenishing enteric neurons continuously lost to apoptosis (Kulkarni et al., 2017). Although fundamental tenets of this proposition are not supported by available experimental evidence (Joseph et al., 2011; Laranjeira et al., 2011; White et al., 2018), it highlights critical but unresolved questions regarding the cellular and molecular mechanisms underpinning the maintenance and regenerative potential of the ENS in vertebrates.

To address these questions, we investigated the ENS of zebrafish, an excellent model organism for studies on NSCs and neural regeneration in vertebrates. Using genetic lineage tracing, gene expression profiling, correlative light and electron microscopy (CLEM), live imaging, and computational modelling, we demonstrate that the non-neuronal compartment of the zebrafish ENS expresses the transgenic reporter *Tg(her4.3:EGFP)* and shares properties with mammalian EGCs and brain NSCs. *Tg(her4.3:EGFP)*^+^ ENS cells exhibit morphological features and express genes characteristic of mammalian enteric glia, but canonical glial markers are undetectable. More akin to radial glial cells (RGCs) of zebrafish brain, EGFP^+^ ENS cells proliferate and undergo constitutive neuronal differentiation which is under the control of Notch signalling. Together, our studies demonstrate the *in vivo* neurogenic potential of enteric glia in vertebrates and reveal previously unanticipated similarities to NSCs in the brain.

## Results

### Expression of canonical glia markers is undetectable in the zebrafish ENS

To pave the way for a systematic search for cells harbouring neurogenic potential in the ENS of non-amniotic vertebrates, we first set out to characterise the non-neuronal compartment of the zebrafish ENS, the most likely source of enteric neural progenitors. Initially, we combined the *SAGFF234A* Gal4 transcriptional activator gene trap with the *UAS:GFP* transgene in order to generate *SAGFF234A;UAS:GFP* animals in which ENS progenitors and their descendants were labelled with GFP (Heanue et al., 2016a; Kawakami et al., 2010). In 7 day post fertilisation (dpf) larvae the majority of GFP^+^ cells (93.76% ± 2.99) co-expressed the pan-neuronal marker HuC/D (Suppl. Fig. 1A-C), suggesting that in comparison to mammals, in which EGCs outnumber enteric neurons (Gabella, 1981; Ruhl, 2005), the non-neuronal ENS cell population of zebrafish is considerably smaller. To support this supposition, we also quantified the proportion of neurons within the ENS of *Tg(−4725sox10:Cre;*β*actin-LoxP-STOP-LoxP-hmgb1-mCherry)* transgenic fish (hereafter abbreviated as *Tg(sox10:Cre;Cherry)*) in which *sox10*-driven Cre recombinase activates a nuclear Cherry reporter in early NC cells and all derivative lineages, including the ENS (Rodrigues et al., 2012; Wang et al., 2011). Consistent with the analysis of *SAGFF234A;UAS:GFP* animals, the majority of Cherry^+^ cells (84.79±7.70%) in the gut of 7 dpf *Tg(sox10:Cre;Cherry)* larvae were positive for HuC/D (Fig. 1A, C). Similar analysis in adult (≥ 3 months old) *Tg(sox10:Cre;Cherry)* zebrafish showed that, although the fraction of non-neuronal Cherry^+^ cells was higher relative to 7 dpf larvae, even at this stage the majority of ENS^+^ cells (65.49±4.8%) were neurons (Fig. 1B, C). Therefore, the non-neuronal compartment in the zebrafish ENS is notably smaller relative to its mammalian counterpart.

**Figure 1.**
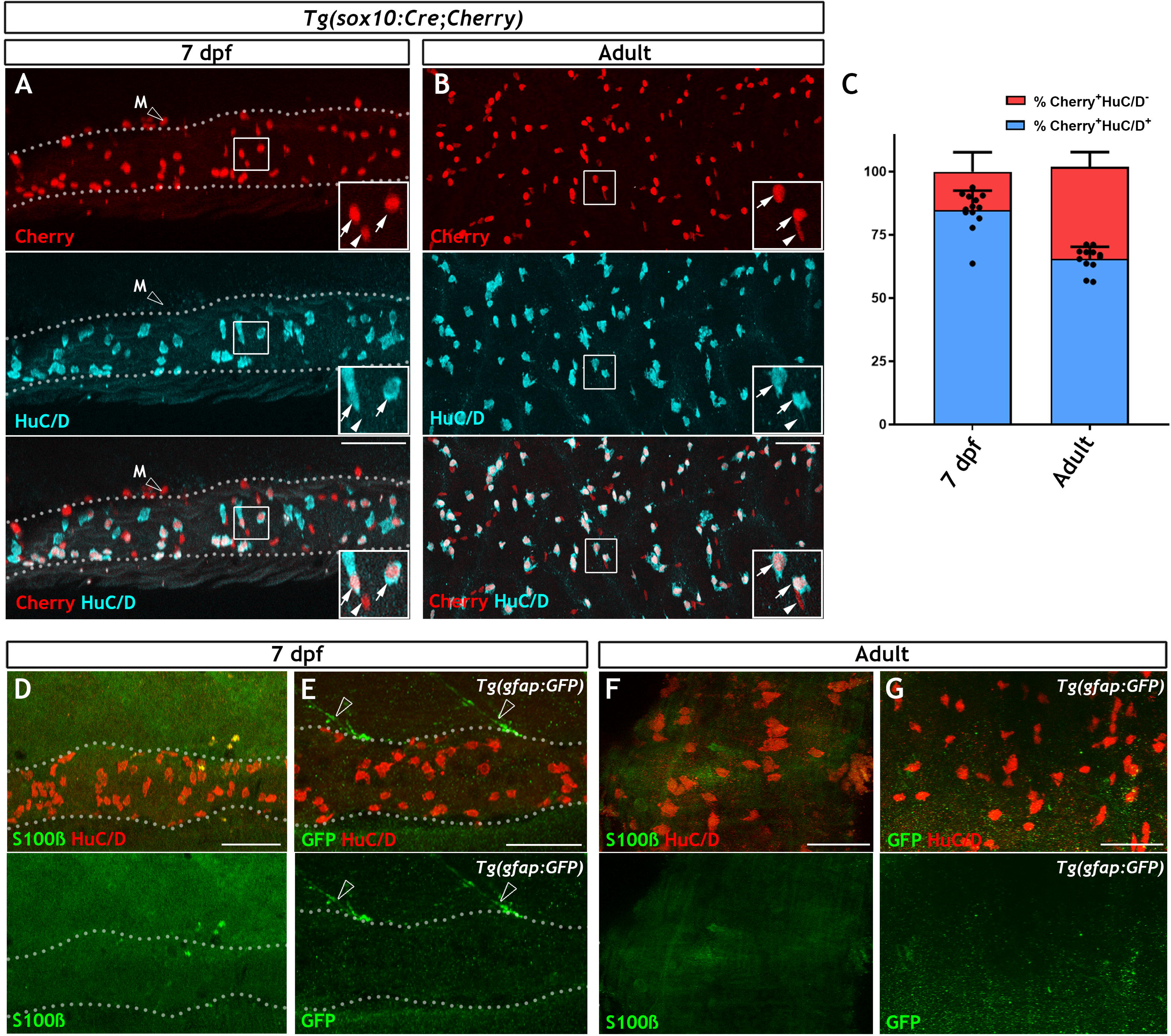
The non-neuronal compartment of the zebrafish ENS is relatively small and is not identified using canonical glial markers. (**A**) Confocal images of the gut of 7dpf *Tg(sox10:Cre;Cherry)* larvae immunostained for Cherry (red, top) and HuC/D (cyan, middle). The bottom panel is a merge of the Cherry and HuC/D signals. Inset shows a high magnification of the boxed area. Arrows point to Cherry^+^HuC/D^+^ cells and arrowhead points to a Cherry^+^HuC/D^-^ cell. Dotted line delineates the gut. Open arrowhead indicates a Cherry^+^ NC-derived melanocyte (M) which is present outside the intestine. (**B**) Confocal images of the ENS in adult zebrafish intestine immunostained for Cherry (red, top) and HuC/D (cyan, middle). The bottom panel is a merge of the Cherry and HuC/D signals. Inset shows a high magnification of the boxed area. Arrows point to Cherry^+^HuC/D^+^ cells and arrowhead points to a Cherry^+^HuC/D^-^ cell. (**C**) Quantification of the neuronal (Cherry^+^HuC/D^+^) and non-neuronal (Cherry^+^HuC/D^-^) cellular compartments within the *sox10*-lineage at 7dpf and adult zebrafish. (**D**) Confocal images of the gut of 7dpf zebrafish larvae immunostained for S100β (green) and HuC/D (red). No s100β signal was detected in the ENS, despite abundant neurons throughout the intestine. (**E**) Confocal images of the gut of 7dpf *Tg(gfap:GFP)* larvae immunostained for GFP (green) and HuC/D (red). No GFP signal was visible within the intestine despite abundant HuC/D^+^ neurons. GFP^+^ fibres associated with spinal nerves are observed descending towards the gut but never enter the intestine (open arrowheads). Dotted lines in D and E delineate the gut. (**F**) Immunostaining of the ENS of adult zebrafish with S100β (green) and HuC/D (red). (**G**) Immunostaining of the ENS of adult *Tg(gfap:GFP)* zebrafish with GFP (green) and HuC/D (red). S100β (F) and GFP (G) signal was absent despite the presence of HuC/D^+^ neurons. 50µm scale bars shown in merge panels.

All non-neuronal cells of the mammalian ENS are identified as enteric glia expressing combinations of the canonical glia markers S100β, GFAP and BFABP (Hao et al., 2016; Young et al., 2003). To determine whether these marker proteins are also expressed in the zebrafish ENS, we used antibodies raised against them to immunostain 7 dpf larvae, a stage when organised intestinal motility patterns controlled by gut-intrinsic neural networks are clearly evident (Heanue et al., 2016a; Holmberg et al., 2007; Kuhlman and Eisen, 2007). Surprisingly, no signal was detected in the ENS of zebrafish at this stage (Fig. 1D and Suppl. Fig. 1D-E). Immunostaining signal detected with two antibodies specific for zebrafish GFAP (Baker et al., 2019; Trevarrow et al., 1990) was likely to represent cross-reactivity with non-neuroectodermal gut tissues as it persisted in *ret* mutant larvae, which lack enteric neuroglia networks (Suppl. Fig. 1F-I) (Heanue et al., 2016a). Immunostaining signal for GFAP has previously reported in the ENS (Baker et al., 2019; Kelsh and Eisen, 2000), however in our experiments the expression is not apparently within the NC-derived lineages. Consistent with the immunostaining, expression of the *Tg(gfap:GFP)* transgene (Bernardos and Raymond, 2006) was also undetectable in the gut of 7dpf larvae (Fig. 1E). In contrast to the ENS, these immunostaining reagents identified the expected signal in the spinal cord (Suppl. Fig. J-O). To ascertain that the lack of glia marker expression was not due to delayed maturation of enteric glia, we also immunostained adult zebrafish gut for GFAP, S100β BFABP and (in the case of *gfap:GFP* transgenics) GFP. Similar to 7dpf animals, no apparent ENS-specific expression of these markers or the *gfap:GFP* transgene detected in adult gut (Fig. 1F-G, Suppl. Fig 1P-Q). Finally, contrary to reports indicating expression of Nestin in non-neuronal cells of mammalian enteric ganglia (Kulkarni et al., 2017), no expression of the *nestin:GFP* transgene was detected in the ENS of adult zebrafish (Suppl. Fig. 1R). Taken together, our studies demonstrate that the non-neuronal compartment of the zebrafish ENS is considerably smaller relative to its mammalian counterpart and cannot be labelled by immunohistochemical reagents commonly used for the identification of enteric glia.

### Non-neuronal cells of the zebrafish ENS share with mammalian EGCs early NC cell and ENS progenitor markers

To explore further the gene expression profile of the non-neuronal ENS cell compartment in zebrafish, we carried out bulk RNA sequencing of fluorescent-labelled nuclei (nRNAseq) isolated from *Tg(sox10:Cre;Cherry)* adult gut muscularis externa. This strategy, which we described recently (Obata et al., 2020), avoids lengthy protocols of tissue dissociation and cell isolation that are often associated with considerable cell damage. Since the available transgenic tools did not allow us to label specifically the non-neuronal ENS cell compartment, bulk nRNAseq was performed on nuclei purified by FACS (fluorescent-activated cell sorting) representing both the Cherry^+^ (entire ENS) and Cherry^-^ (non-ENS) muscularis externa cell populations of *Tg(sox10:Cre;Cherry)* zebrafish gut (Fig. 2A and Suppl. Fig. 2A; see also Materials and Methods). Principal component analysis (PCA) demonstrated a clear separation of the Cherry^+^ and Cherry^-^ nuclear transcriptomes along PC1 (Suppl. Fig. 2B), indicating that variability along this axis is determined predominantly by the lineage origin (NC vs non-NC) of the two cell populations. As expected, genes associated with non-NC tissues, such as smooth muscle cells (*mylka*, *myh11a*, *cald1a*, *srfa*, *gata6*, *anxa2b*), interstitial cells of Cajal (*ano1*, *kita*, *kitb*) and immune cells (*lcp1*, *lck*, *lyz*), were upregulated in the Cherry^-^ nuclear transcriptome (Fig 2B). Conversely, genes associated with the NC-derived ENS lineages (such as *elavl3*, *elavl4*, *ret*, *vip*, *chata*, *sox10*) were upregulated in the Cherry^+^ nuclear population (Fig 2B, Crick weblink will be made available for interactive data analysis). Furthermore, gene ontology (GO) terms enriched in the Cherry^+^ nuclear population were associated with nervous system development and function (Suppl. Fig 2C-E). Finally, direct comparison of the Cherry^+^ dataset to the transcriptional profile of enteric neurons from 7dpf larvae expressing the *Tg(phox2b:EGFP)^w37^* transgene (Roy-Carson et al., 2017), identified a large cohort of shared genes (including *phox2bb*, *ret*, *elavl3*, *elavl4*, *vip*, *nmu*) that presumably reflect the neural component of the mixed Cherry^+^ nuclear population (Fig 2C, yellow dots, Suppl. Fig 2F and Suppl. Table 1).

**Figure 2.**
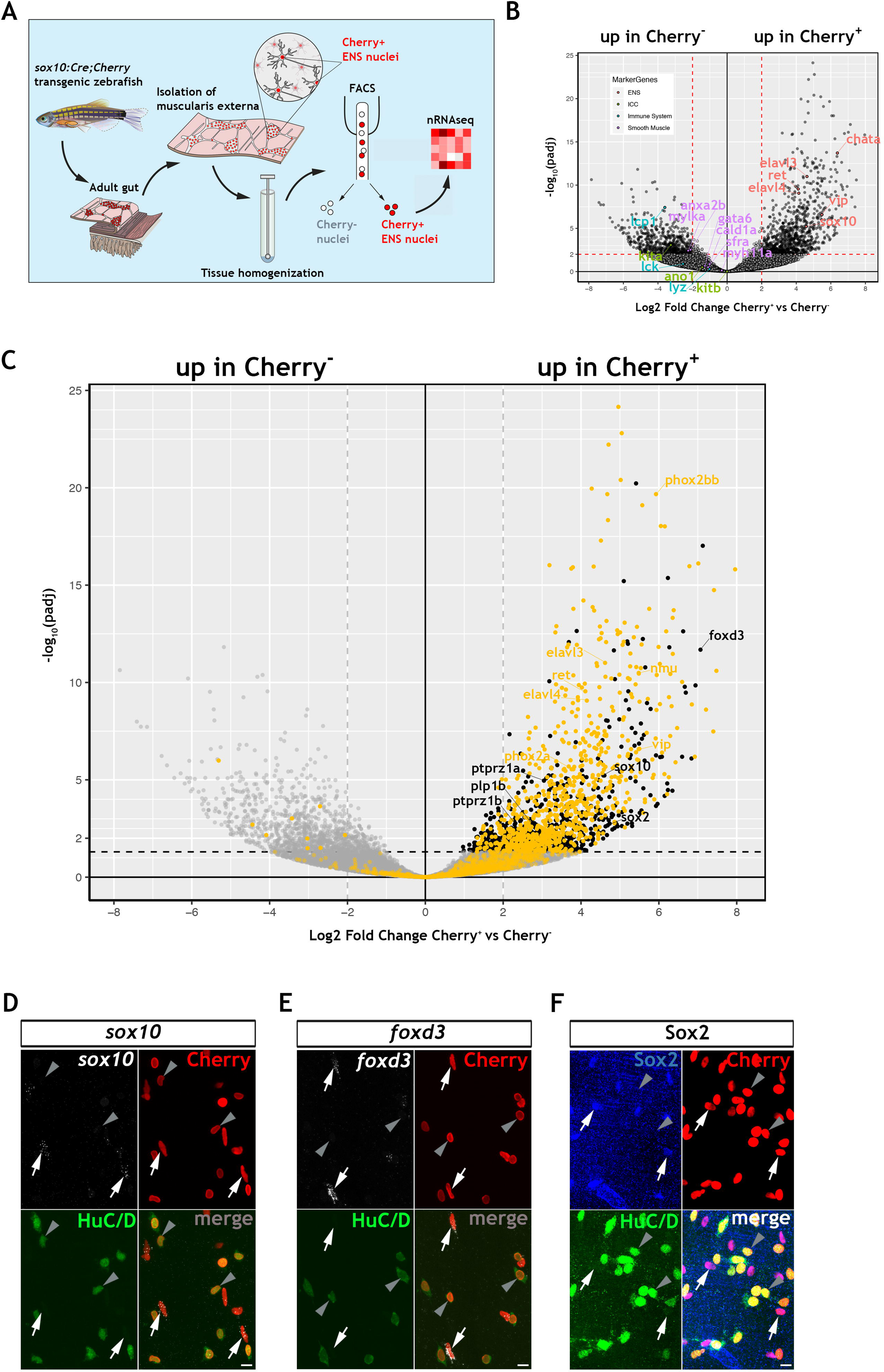
Transcriptomic profiling of the adult zebrafish ENS. (**A**) Experimental strategy for the isolation of ENS nuclei from adult *Tg(sox10:Cre;Cherry)* guts and nuclear RNAseq. (**B**) Volcano plot shows mean log_2_ fold-change (x axis) and significance (-log_10_ adjusted p-value) (y axis) of genes differentially expressed in Cherry^+^ relative to Cherry^-^ nuclei. Genes characteristic of the ENS are highlighted in red and are more abundant in Cherry^+^ nuclei, whereas genes characteristic of non-neuroectodermal lineages, such as smooth muscle (purple), interstitial cells of Cajal (green) and immune associated (blue), are more abundant in Cherry^-^ nuclei. (**C**) Volcano plot (as in B) in which genes previously identified in a transcriptional characterization of larval ENS neurons (Roy-Carson et al., 2017) are shown in yellow. These include established neuronal markers, such as *phox2bb*, *ret*, *elavl3*, *elavl4*, *vip*, and *nmu*. Genes enriched in the Cherry^+^ nuclear population but absent from the larval ENS neuron transcriptome are shown in black. These include *sox10*, *foxd3*, *sox2*, *plp1*, the mammalian orthologues of which are expressed by mouse EGCs, as well as *ptprz1a*, and *ptprz1b*, which have been identified in glioblastoma stem cells. Genes with padj < 0.05 (Log_10_ p-value < 1.3) and/or log_2_FC < 0 are shown in grey. (**D,E**) Fluorescent *in situ* hybridization (RNAscope) using probes for *sox10* (D) and *foxd3* (E) on adult *Tg(sox10:Cre;Cherry)* gut muscularis externa preparations immunostained for Cherry (ENS lineage) and HuC/D (ENS neurons). Signal for both *sox10* and *foxd3* (white arrows) corresponds to non-neuronal cells (Cherry^+^HuC/D^-^, arrows) but was absent from enteric neurons (Cherry^+^HuC/D^+^, arrowheads). (**F**) Immunostaining of adult *Tg(sox10:Cre;Cherry)* gut for Sox2 (blue), Cherry (red) and HuC/D (green). Sox2 is expressed specifically by non-neuronal ENS cells. 10µm scale bars shown in merge panels.

To identify genes expressed by the non-neuronal compartment of the zebrafish ENS we next compared the Cherry^+^ dataset to a recently reported transcriptome of mouse EGCs, which includes a list of the 25 most highly expressed genes in PLP1^+^ enteric glia (Rao et al., 2015). Zebrafish orthologues for several genes in this list were enriched in the Cherry^+^ transcriptome (Suppl. Fig. 2G), suggesting that they are expressed by the non-neuronal cells of the zebrafish ENS. Among these genes were *sox10* and *foxd3*, which in mammals are expressed by early NC cells and ENS progenitors and maintained in enteric glia (Mundell and Labosky, 2011; Mundell et al., 2012; Weider and Wegner, 2017), as well as genes with established association to glial cells, such as *plp1*, *ptprz1a* and *ptprz1b* (Fujikawa et al., 2017). Having delineated the neural component of the Cherry^+^ transcriptome (Fig. 2C, yellow dots, and Suppl. Fig. 2F), we removed this cohort of genes in order to enrich for transcripts of the non-neuronal ENS cell compartment (Fig. 2C, black dots, Suppl. Fig. 2H and Suppl. Table 2). This strategy highlighted several genes that were identified by our previous analysis, including *sox10* and *foxd3*, and numerous additional genes, including *sox2*, which is expressed by mouse ENS progenitors and adult EGCs (Belkind-Gerson et al., 2017; Heanue and Pachnis, 2011; Rao et al., 2015). Expression of *sox10*, *foxd3* and *sox2* in the non-neuronal compartment of the zebrafish ENS was validated by combining multiplex fluorescence *in situ* hybridisation (RNAscope) with immunostaining for HuC/D and the Cherry reporter on muscularis externa preparations from the gut of adult *Tg(sox10:Cre;Cherry)* zebrafish (Fig. 2D-F). Consistent with our immunostaining analysis, which failed to detect canonical glia marker expression in the zebrafish ENS (Fig 1F, G and Suppl. Fig. 1P, Q), transcripts for *gfap*, *s100b* and *fabp7a/b* were also absent from the Cherry^+^ nuclear transcriptome. We cannot exclude the possibility that such markers may be revealed by in depth sequencing of single cells. Together, these experiments indicate that, despite our failure to detect expression of commonly used EGC markers, the transcriptomes of the non-neuronal compartment of the zebrafish ENS and mammalian enteric glia have considerable overlap, including gene associated with early NC cells and ENS progenitors.

### Non-neuronal cells in the adult zebrafish ENS express the Notch activity reporter ***Tg(her4.3:EGFP)***

In mammals, Notch signalling promotes enteric gliogenesis by attenuating a cell-autonomous neurogenic programme of ENS progenitors (Okamura and Saga, 2008), but the expression of Notch target genes in adult EGCs is unclear. To examine whether Notch signalling is active in non-neuronal cells of the zebrafish ENS, we examined adult gut for expression of the transgenic Notch activity reporter *Tg(her4.3:EGFP)* (see Materials and Methods for the nomenclature of this transgene), which marks NSCs and neural progenitors in the brain (Alunni and Bally-Cuif, 2016; Yeo et al., 2007). This analysis identified a network of GFP^+^ cells in the muscularis externa of the gut that was closely associated with enteric neurons and their projections (Fig. 3A and Suppl. Fig. 3A). To provide direct evidence that *Tg(her4.3:EGFP)*-expressing cells are integral to the ENS, we introduced the *her4.3:EGFP* transgene into the *Tg(sox10:Cre;Cherry)* genetic background and immunostained gut preparations from adult *Tg(her4.3:EGFP;sox10:Cre;Cherry)* animals for GFP, HuC/D and Cherry. As expected, GFP^+^ cells were negative for HuC/D but expressed the Cherry reporter (Fig. 3C), indicating that they belong to the non-neuronal compartment of the ENS.

**Figure 3.**
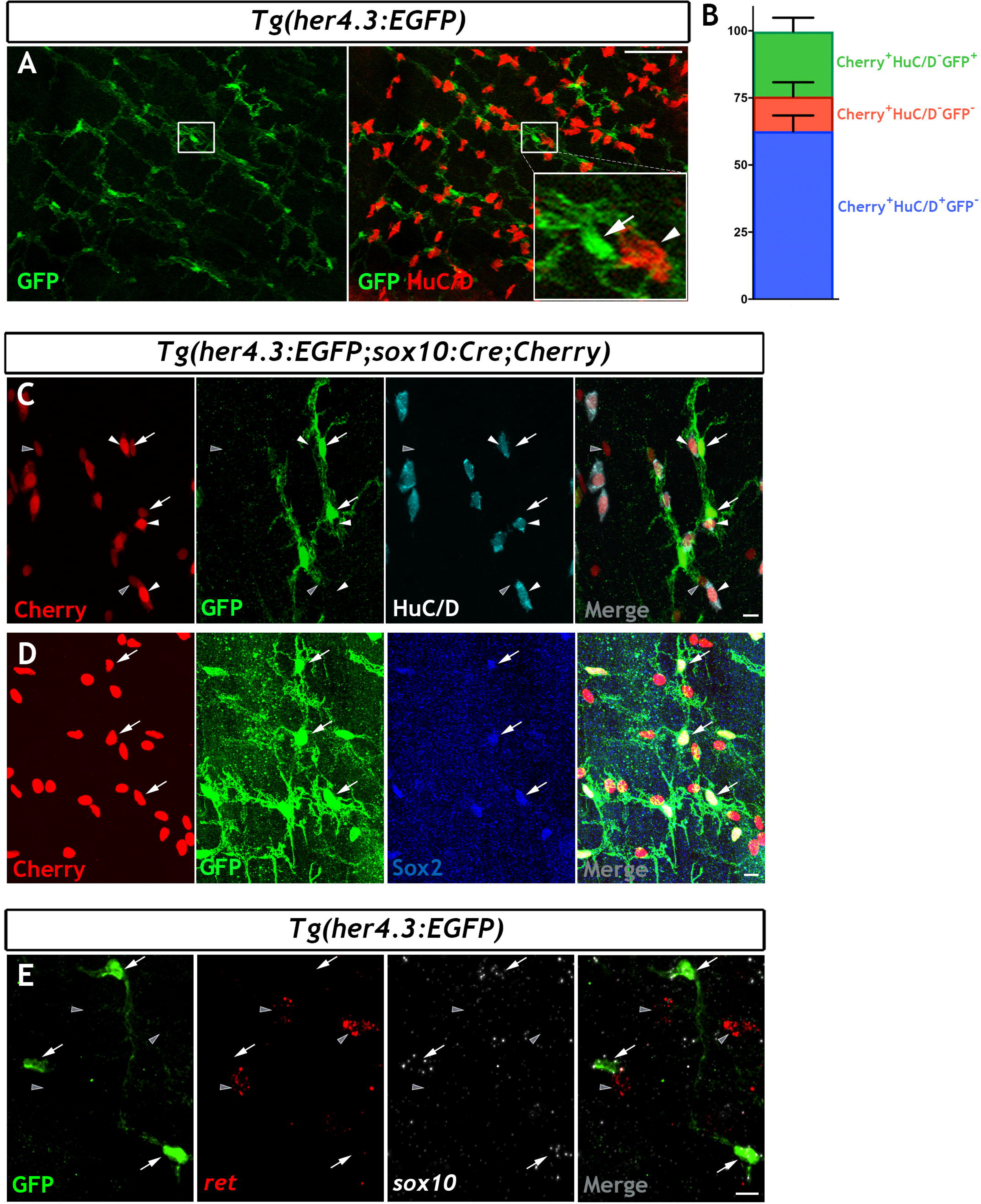
The *her4.3:EGFP* transgene is a novel marker of the non-neuronal cell population in the zebrafish ENS. (**A**) Confocal images of adult *Tg(her4.3:EGFP)* zebrafish gut immunostained for GFP (green) and HuC/D (red). Inset is a high magnification of boxed area showing that GFP^+^ cells (arrow) are closely associated with HuC/D^+^ neurons (arrowhead). (**B**) Quantification of neuronal (Cherry^+^ HuC/D^+^GFP^-^, blue) and non-neuronal cell populations (Cherry^+^HuC/D^-^GFP^+^ and Cherry^+^HuC/D^-^GFP^-^, green and red, respectively) in the ENS of adult *Tg(her4.3:EGFP*;*sox10:Cre;Cherry)* zebrafish. (**C**) Confocal images of the ENS from *Tg(her4.3:EGFP*;*sox10:Cre;Cherry)* zebrafish immunostained for Cherry (red), GFP (green) and HuC/D (cyan). Note the presence of Cherry^+^HuC/D^-^GFP^+^ (arrows) and Cherry^+^ HuC/D^-^ GFP^-^ (grey arrowheads) cells as well as the presence of Cherry^+^HuC/D^+^GFP^-^ neurons (white arrowheads). (**D**) Immunostaining of adult *Tg(her4.3:EGFP*;*sox10:Cre;Cherry)* gut with antibodies for Cherry (red), GFP (green) and Sox2 (blue). Arrows point to cells expressing all three markers. (**E**) RNAscope analysis for *ret* and *sox10* on ENS preparations from *Tg(her4.3:EGFP*) zebrafish guts immunostained for GFP. Note that GFP^+^ cells (arrows) express *sox10* and are found in proximity to *ret*^+^GFP^-^ enteric neurons (grey arrowheads). Scale bars in merge panels: (A) 50µm (C-E) 10µm.

Consistent with this idea, GFP^+^ cells co-expressed *sox2* and *sox10* (Fig. 3D, E), which were identified by our transcriptomic analysis as genes expressed by the non-neuronal compartment of the zebrafish ENS. The GFP^+^HuC/D^-^ cell population in *Tg(her4.3:EGFP;sox10:Cre;Cherry)* represented approximately a quarter (24.20 ± 5.18%) of all Cherry^+^ ENS cells, but 12.93 ± 5.33% of Cherry^+^ cells were negative for both GFP and HuC/D (Cherry^+^GFP^-^HuC/D^-^) (Fig. 3B). Therefore, the majority of non-neuronal ENS cells in adult zebrafish gut can be identified by the expression of the Notch activity reporter *Tg(her4.3:EGFP)*.

### GFP^+^ cells in the ENS of *Tg(her4.3:EGFP)* zebrafish have morphological characteristics of mammalian EGCs

To provide evidence that *Tg(her4.3:EGFP)*-expressing cells in the zebrafish ENS are equivalent to mammalian EGCs, we characterised the morphology of GFP^+^ cells in the gut of *Tg(her4.3:EGFP)* transgenics. At the light microscopy level GFP^+^ cells were highly branched and formed four morphological groups that generally corresponded to the four morphological subtypes of mouse EGCs (Suppl. Fig. 3B-E) (Boesmans et al., 2015; Gulbransen and Sharkey, 2012). In addition to the muscularis externa (Suppl Fig 3B, C, E), GFP^+^ cells were also found within the mucosa in close proximity to the intestinal epithelium (Suppl Fig. 3D), similar to type III mucosal EGCs located within the lamina propria of the mammalian gut (Boesmans et al., 2015; Kabouridis et al., 2015).

Mammalian EGCs have unique ultrastructural features and establish characteristic contacts with enteric neurons and their projections (Gabella, 1972, 1981). To determine whether similar features are exhibited by the GFP^+^ ENS cell population in *Tg(her4.3:EGFP)* zebrafish, we analysed EGFP^+^ cells in *Tg(her4.3:EGFP;SAGFF217B;UAS:mmCherry)* transgenics using CLEM (Muller-Reichert and Verkade, 2012). In these animals, EGFP marks non-neuronal ENS cells while Cherry, which is driven by the binary reporter *Tg(SAGFF217B;UAS:mmCherry)* (Kawakami et al., 2010), labels a subset of enteric neurons (Suppl. Fig. 4A). CLEM confirmed the close association of EGFP^+^ cells with enteric neurons and their projections (Fig. 4, Suppl. Fig. 4B,C and Supplementary Movie 1). Processes emanating from EGFP^+^ cells directly contacted enteric neurons (Fig. 4B, D and Suppl. Fig. 4C), but similar to mammalian EGCs (Gabella, 1981) they did not form complete “capsules” around neuronal somata, allowing large parts of enteric neurons to be in direct contact with adjacent cells (Fig. 4A, B, Suppl. Fig. 4C and Supplementary Movie 1). EGFP^+^ cells also extended complex sheet-like extensions, which frequently enclosed and/or subdivided the tightly packed bundles of neural projections into sectors (Fig. 4D, Suppl. Fig. 4C and Supplementary Movie 1). Deep nuclear crenations, a characteristic feature of mammalian EGCs and other populations of peripheral glial cells (Gabella, 1981), were also found in the nuclei of EGFP^+^ cells (Fig. 4B, D and Suppl. Fig. 4C). Together, our gene expression and morphological analysis argues that, despite the lack of canonical glia marker expression, the cell population expressing the Notch activity reporter *Tg(her4.3:EGFP)* corresponds to mammalian EGCs. Henceforth, we will be referring to *Tg(her4.3:EGFP)*-expressing cells in the adult zebrafish ENS as EGCs.

**Figure 4.**
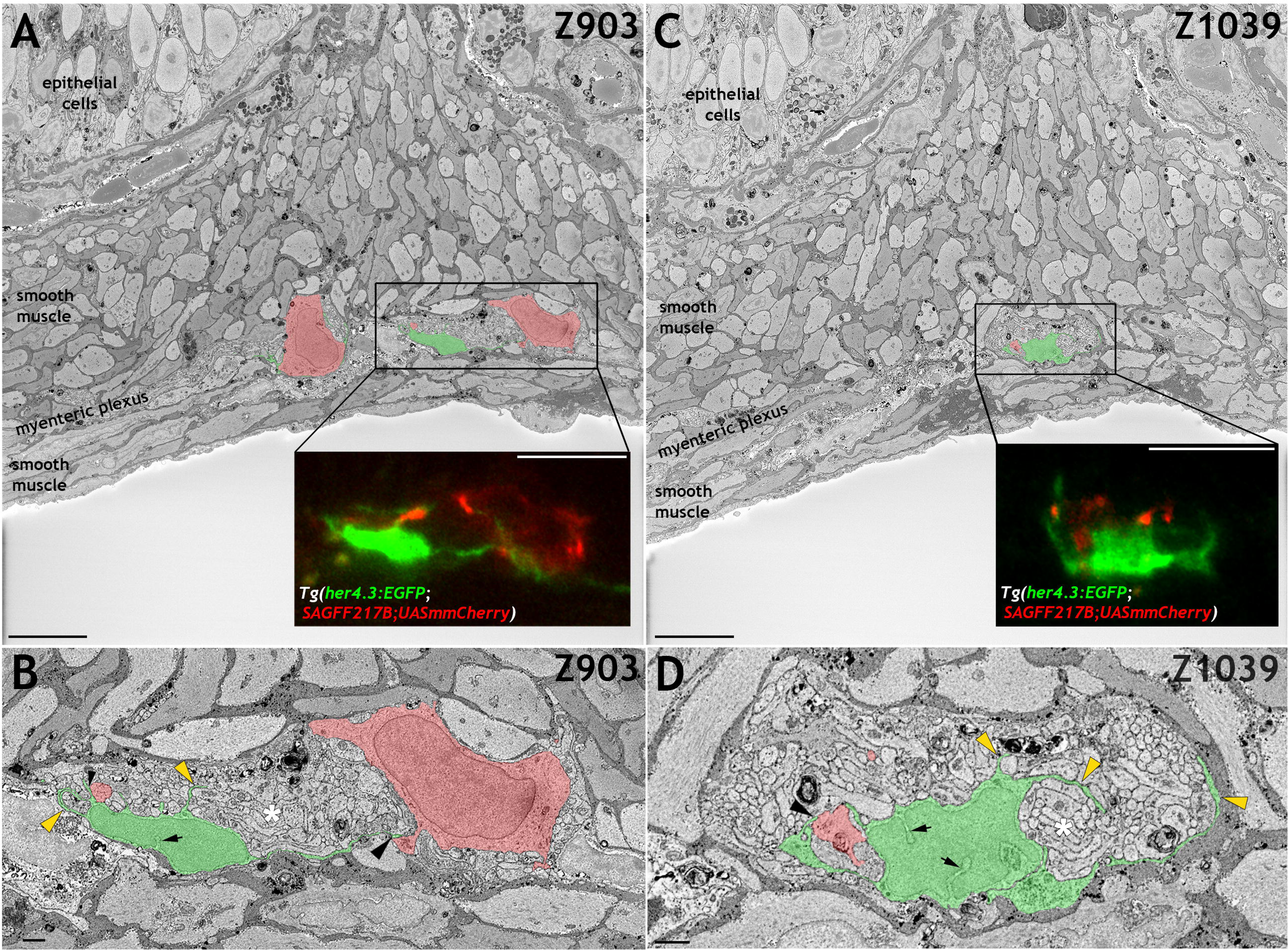
*her4.3:EGFP-*expressing cells in the zebrafish ENS share with mammalian enteric glia characteristic ultrastructural features. (**A** and **C**) Electron micrographs (z-stack # z903 in A and #1039 in C) from a 3D region of interest from the midgut of *Tg(her4.3:EGFP;SAGFF217;UAS:mmCherry)* zebrafish. Insets shows super-resolution light microscopy images of EGFP^+^ non-neuronal cells and mmCherry^+^ neurons that correspond to the boxed areas of the electron micrograph. (**B** and **D**) High-resolution images of the boxed areas shown in A (B) and C (D). GFP^+^ cells are pseudocoloured in green and enteric neurons in red. Black arrowheads indicate points of contact between GFP^+^ processes and neurons. Yellow arrowheads indicate GFP^+^ sheet-like extensions that compartmentalise axon bundles (white asterisks). Nuclear crenelations in nuclei of GFP^+^ cells are indicated with black arrows. Scale bars: 10µm (A, C and insets A,C) and 1µm (B,D).

### Developmental profile of zebrafish EGCs

To examine the developmental profile of zebrafish EGCs, we immunostained *Tg(her4.3:EGFP;SAGFF234A;UAS:mmCherry)* transgenics for GFP and Cherry at different developmental stages. At 54 hours post fertilisation (hpf), a stage at which NC cell-derived Cherry^+^ cells are restricted to two distinct migratory columns along the gut (Heanue et al., 2016), no double positive (Cherry^+^GFP^+^) cells were identified (Suppl. Fig. 5A). However, at 60hpf a small number of GFP^+^ cells were discernible within the Cherry^+^ streams of NC cells (Suppl. Fig. 5B) and became more abundant in 4dpf larvae (Suppl. Fig. 5C). To further examine the developmental dynamics of the GFP^+^ cell lineage, we performed time-lapse confocal microscopy of live *Tg(her4.3:EGFP;SAGFF234A;UAS:mmcherry)* embryos at similar stages. Imaging commenced at 56 hpf with the migratory front of mmCherry^+^ NC cell columns positioned at the rostral end of the field of view (Heanue et al., 2016a) and continued for 40 hours (1 image every 10 minutes). Consistent with the analysis performed on fixed embryos, no EGFP^+^ cells were identified within the mmCherry^+^ population during the first hours of imaging (Fig. 5A). However, EGFP^+^ cells appeared within the columns of mmCherry^+^ cells at around 62hpf (Fig. 5B), more than 90µm behind the front of migrating mmCherry^+^ NC cells, and the number of GFP^+^ cells increased over the remaining imaging period (Fig. 5C, D) (Supplementary Movie 2). On several occasions, we identified individual mmCherry^+^ cells inducing *de novo* expression of EGFP (Supplementary Movie 3). EGFP^+^ cells emerged in a rostro-caudal sequence mirroring the wave of ENS neuron maturation (Heanue et al., 2016b) but they were almost always located behind the front of migrating enteric NC cells. Relative to the tip of the mmCherry^+^ migratory column, which was displaced caudally at a constant rate until it reached the caudal end of the FOV, EGFP^+^ cells on average exhibited minimal rostrocaudal displacement (Fig. 5E; 132 EGFP^+^ cells analysed from 4 fish), suggesting that during ENS development the *her4.3:EGFP* transgene is expressed in post-migratory cells.

**Figure 5.**
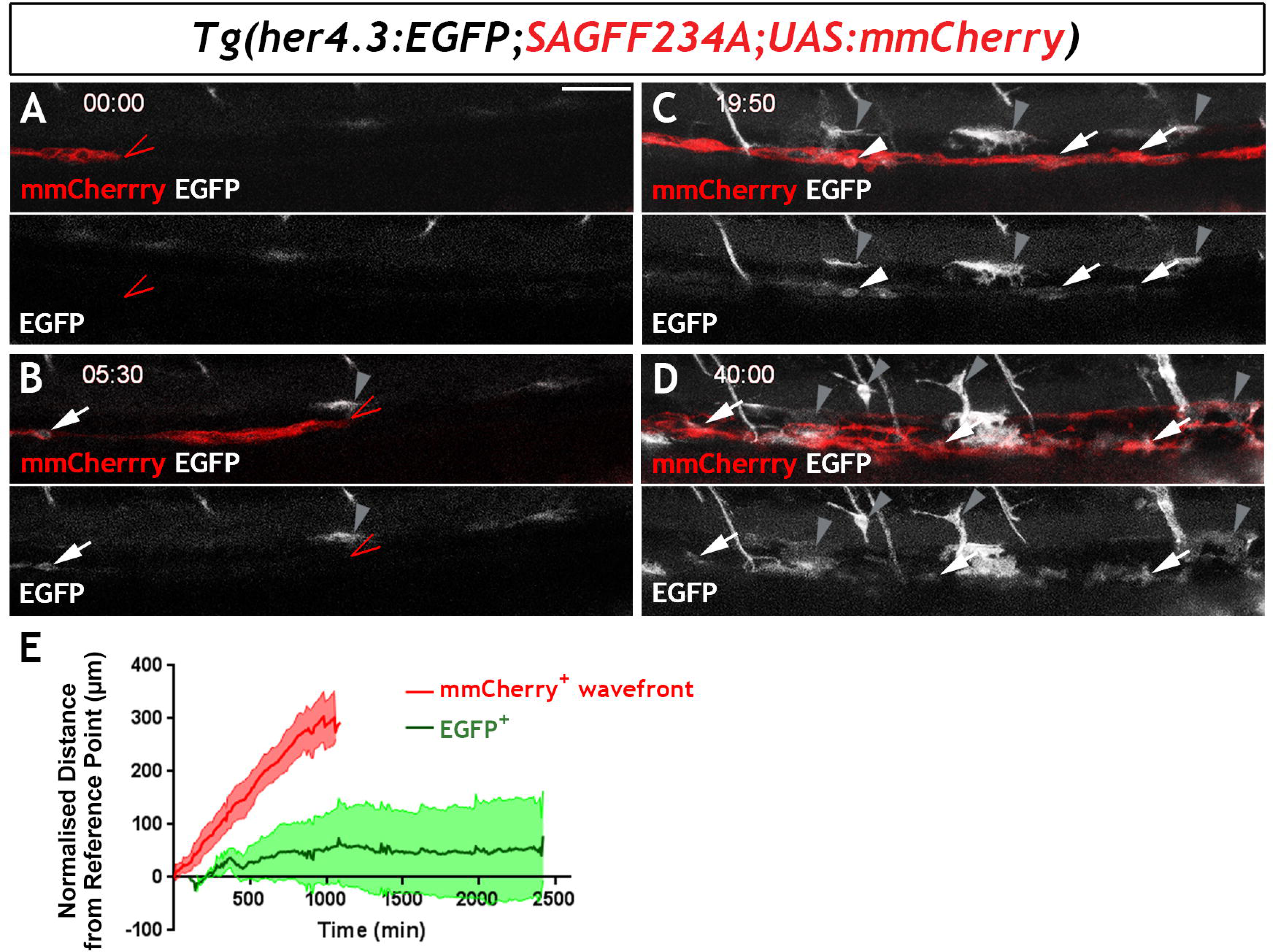
Live imaging of *Tg(her4.3:EGFP)^+^* cell ontogenesis in the zebrafish ENS. (**A-D**) Still images from time-lapse recording of a *Tg(her4.3:EGFP;SAGFF234A;UAS:mmCherry)* embryo imaged from 56hpf (denoted as 00:00) until 96hpf (40:00). At 00:00 (A) the mmCherry^+^ wavefront of NC cells (red, red arrowhead) is at the rostral side of the field of view (FOV) and no EGFP^+^ cells (grey) are present. At 05:30 (B), the first EGFP^+^ cells (grey, arrow) appear within the mmCherry^+^ NC cell column (red), behind the migratory wavefront. Bright GFP^+^ melanocytes are designated (grey arrowheads) . (C) At 19:50 the NC cell column extends throughout the FOV and the number of EGFP^+^ cells (grey, arrows) has increased. Arrowhead points to an EGFP^+^ cell exhibiting a rounded morphology, which can be seen to divide in subsequent time lapse images. An increasing number of bright GFP^+^ melanocytes appear (grey arrowheads), and are relatively static in the time lapse movies. (D) At the end of the recording (40:00), EGFP^+^ cells (grey) can be found throughout the gut (white arrowheads). Abundant brightly GFP^+^ melanocytes are present in the gut region (grey arrowheads), whose characteristic morphology is apparent. (**E**) Quantification of cell displacement (normalised distance from reference point/time) of the mmCherry^+^ wavefront (red) and EGFP^+^ cells (green). Data are given as mean ± SD. 50µm scale bar in A.

Next, we characterised the cell division patterns of the 79 EGFP^+^ cells that migrated into the field of view or arose *de novo* during the live imaging period. Of these, 37 cells gave rise to at least one generation of GFP^+^ progeny. 26 cells (∼33%) underwent a single cell division generating two daughters, many of which lost EGFP expression over the course of imaging. In these cases the EGFP expressing cells were not migratory and the EGFP expression diminished and then extinguished. In a proportion of cells (8 cells; ∼10%), after a first division event, one or both of the daughter cells underwent a further cell division, generating EGFP^+^ granddaughters, some of which lost expression of the reporter. And for 3 cells (∼4%), following two division events, one granddaughter cell underwent a further division to generate a third generation of EGFP^+^ progeny. Altogether, 53 EGFP^+^ cells were seen to undergo a cell division event during the imaging period. Therefore, during development *Tg(her4.3:EGFP)*-expressing cells are capable of entering the cell cycle but those that do so undergo only a limited number of cell divisions and many of their progeny eventually lose expression of EGFP. Loss of EGFP signal could be associated with neuronal differentiation since we occasionally identified in the gut of 7 dpf *her4.3:EGFP* transgenic larvae cells that were weakly immunostained for both HuC/D and GFP (Suppl Fig 5D). Taken together, our analysis of *Tg(her4.3:EGFP)* expression during zebrafish development suggests that nascent EGCs are postmigratory NC-derived cells which maintain proliferative and neurogenic potential.

### Proliferation and neuronal differentiation of zebrafish EGCs during homeostasis

Enteric glia in adult mammals are generally quiescent with only a small fraction of cells undergoing cell division at any given time (Joseph et al., 2011). To examine the proliferative potential of EGCs in adult zebrafish, we immunostained whole-mount gut preparations from adult *Tg(her4.3:EGFP)* transgenics for the proliferation marker mini-chromosome maintenance 5 (MCM5) (Ryu et al., 2005). 10.8±4.2% of GFP^+^ cells were positive for MCM5 (Suppl. Fig. 6), indicating that in contrast to their mammalian counterparts, a considerable proportion of zebrafish EGCs are cycling during homeostasis.

Our earlier observation that EGFP^+^ cells in the ENS of *Tg(her4.3:EGFP)* zebrafish embryos undergo only a limited number of cell divisions suggested that EdU (5-ethynyl-2’-deoxyuridine) labelling of EGCs in adult animals could be used to trace the progeny of proliferating cells and determine their fate. Consistent with the MCM5 immunostaining, we found that at the end of a 3-day EdU labelling pulse (t0), 8.0±4.3% of GFP^+^ cells in the gut of 3 month old *her4.3:EGFP* transgenic zebrafish were co-labelled with EdU (Fig. 6A, B and D). At this stage the majority of GFP^+^EdU^+^ cells formed doublets composed of cells with similar morphology and GFP signal intensity (Fig. 6B and Suppl. Fig. 7B). Occasionally, one or both cells in the doublets exhibited reduced GFP signal (Suppl. Fig. 7C and D), suggesting that, similar to larval stages, the daughters of dividing EGCs in adult *her4.3:EGFP* transgenic zebrafish differentiate into GFP^-^ enteric neurons. This idea was supported by the identification 4 days post-labelling (t4 chase) of EdU^+^ doublets that included GFP^+^HuC/D^-^ and GFP^-^HuC/D^+^cells (Fig. 6A, C). The loss of GFP signal from the daughters of proliferating EGCs cells was also supported by cell population analysis which demonstrated a reduction in the percentage of EdU^+^GFP^+^ cells (t4: 3.6±3.4%, p=6.01×10^-7^; t11: 3.9±3.8%, p=7.61×10^-6^) (Fig. 6D). Interestingly, the reduced percentage of EdU^+^GFP^+^ cells during the EdU chase period was associated with a concomitant increase in the representation of EdU^+^ enteric neurons at t4 (0.71±0.80%, p=6.0×10^-7^) and t11 (0.70±0.82%, p=1.5×10^-6^) relative to t0 (0.068±0.13%) (Fig. 6E). Together, these experiments suggest that the progeny of proliferating EGCs in the zebrafish ENS can differentiate into neurons under physiological conditions.

**Figure 6.**
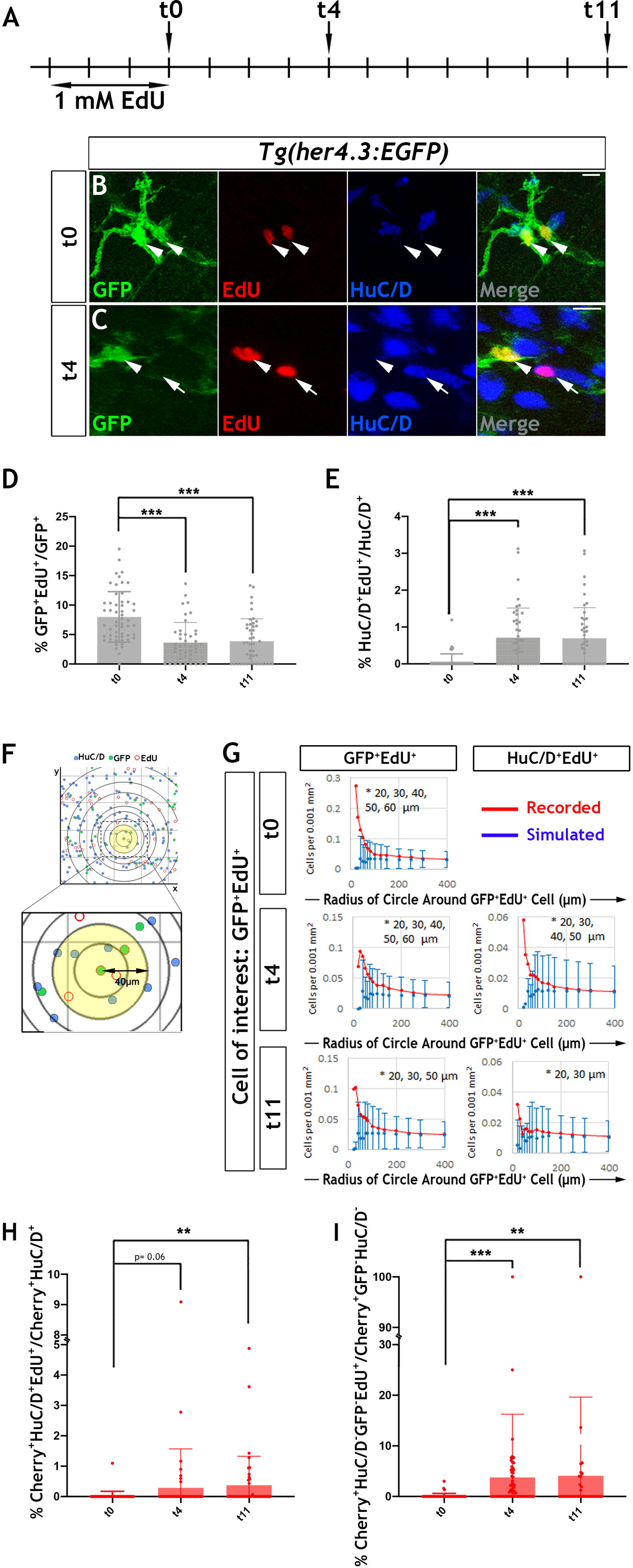
Proliferation and neurogenic differentiation of her4.3:EGFP^+^ ENS cells during homeostasis. (**A**) Schematic representation of experimental design. Adult *Tg(her4.3:EGFP)* zebrafish were immersed in 1mM EdU for three days and analysed at 0 (t0), 4 (t4) or 11 (t11) days after EdU pulse. (**B-C**) GFP (green) and HuC/D (blue) immunostaining of intestines from EdU (red) pulsed animals harvested at t0 (B) and t4 (C). Arrowheads (in B and C) point to GFP^+^HuC/D^-^EdU^+^ cells. Arrow (in C) indicates a GFP^-^ HuC/D^+^EdU^+^ neuron. 10 µm scale bars in B-C merge panels. (**D**) Quantification of the percentage of GFP^+^ cells labelled with EdU at t0, t4 and t1 (mean ±SD). (**E**) Quantification of the percentage of EdU-labelled enteric neurons at t0, t4 and t11 (mean ±SD.) (**F**) Strategy for computational analyses of the density of EdU-labelled HuC/D^+^ and EGFP^+^ cells. EdU^+^GFP^+^ cells were positioned at the centre of concentric circles of increasing radius and the density of EdU^+^GFP^+^ and EdU^+^HuC/D^+^ cells within each circle was calculated. An example of a 40 μm radius circle (yellow) is shown in higher magnification. (**G**) Recorded (red graph) and simulated (blue graph) densities of EdU^+^HuC/D^+^ and EdU^+^GFP^+^ cells (y axis) in concentric circles of increasing radius (x axis) around EdU^+^GFP^+^ cells. Monte Carlo simulation of random distribution of EdU^+^HuC/D^+^ or EdU^+^GFP^+^ cells were performed >2000 times for each dataset in order to establish baseline densities arising in randomly mixed populations. Error bars represent mean ± 90% confidence intervals. At all time-points analysed, recorded densities of EdU^+^HuC/D^+^ and EdU^+^GFP^+^ cells were above the confidence interval (bars) of the simulated densities in 20µm and 60 μm circles (indicated by asterisk). (**H**, **I**) Quantification of the percentage of EdU-labelled Cherry^+^HuC/D^+^ neurons (H) and Cherry^+^GFP^-^HuC/D^-^ cells (I) at t0, t4 and t11 in the intestine of *her4.3:gfp;sox10:Cre;Cherry* transgenics pulse-labelled with EdU according to the protocol shown in panel A. In D-E and H-I, * P<0.05, ** P<0.01, *** P<0.001.

To provide further evidence in support of the lineage relationship between GFP^+^EdU^+^ cells and newborn enteric neurons (HuC/D^+^EdU^+^), we used confocal microscopy and mathematical modelling to estimate the densities of these cell types within circles of increasing radius centred on EdU^+^ cells (Fig. 6F) (Tay et al., 2017). We reasoned that closer proximity of HuC/D^+^EdU^+^ and GFP^+^EdU^+^ cells relative to that expected from random distribution of lineally unrelated cells would indicate origin from common progenitors undergoing cell division. The densities observed at t0, t4 and t11 were compared to values of uniformly distributed cell types generated randomly by Monte Carlo simulations (>2×10^3^ per sampling time). This analysis revealed that the actual densities of GFP^+^EdU^+^ and HuC/D^+^ EdU^+^ cells were significantly higher within the smaller radius circles (<60 µm from the cell of interest) in comparison to those expected by chance, suggesting that the observed homotypic (GFP^+^EdU^+^/GFP^+^EdU^+^) and heterotypic (GFP^+^EdU^+^/HuC/D^+^EdU^+^) clusters of EdU^+^ ENS cells were descendants of a common proliferating progenitor (Fig. 6G). EdU^-^ cells exhibited densities similar to those expected in randomly mixed populations (data not shown). This analysis provides further support to the idea that descendants of proliferating *Tg(her4.3:EGFP)*-expressing ENS cells are capable of undergoing neuronal differentiation in the gut of adult zebrafish.

Next, we considered the possibility that the GFP^-^ non-neuronal ENS cell population (Fig. 3B) is also derived from GFP^+^ progenitors and represents an intermediate stage of neurogenic commitment, in a process analogous to the differentiation of GFP^+^ RGCs in the pallium of *her4.3:EGFP* transgenic zebrafish. To examine this, we pulse-labelled 3 month old *Tg(her4.3:EGFP;sox10:Cre;Cherry)* transgenics with EdU (Fig. 6A) and followed the descendants of proliferating EGCs in the context of the entire ENS lineage. Consistent with our previous analysis (Fig. 6E), the percentage of enteric neurons labelled by EdU (Cherry^+^HuC/D^+^EdU^+^) at t4 and t11 was higher relative to t0 (t0: 0.021±0.15%; t4: 0.28±1.2%, p=0.06; t11: 0.37±0.95%, p=0.0014) (Fig. 6H). Interestingly, this increase was paralleled by an increased percentage of EdU-labelled GFP^-^ non-neuronal ENS cells (Cherry^+^GFP^-^HuC/D^-^EdU^+^) at t4 and t11, relative to t0 (t0: 0.12±0.5%; t4: 3.7±12.5%, p=1.84×10^-6^; t11: 4.1±15.5%, p=0.0024) (Fig. 6I). Together these studies suggest that loss of *Tg(her4.3:EGFP)* expression in the daughters of proliferating EGCs is likely to reflect neurogenic commitment preceding overt neuronal differentiation.

### Notch signalling regulates neuronal differentiation

Inhibition of Notch signalling promotes the proliferation and neurogenic differentiation of *Tg(her4.3:EGFP)*-expressing RGCs in the telencephalon of zebrafish (Alunni et al., 2013; Chapouton et al., 2010). This, together with the observed downregulation of the *her4.3:EGFP* transgene upon neuronal differentiation of GFP^+^ cells (Fig. 6C), suggested that canonical Notch activity regulates the proliferation and differentiation dynamics of EGCs in zebrafish. To examine this possibility, we blocked Notch signalling in adult zebrafish by treating them with the γ-secretase inhibitor LY411575 (referred to as LY) (Alunni et al., 2013; Rothenaigner et al., 2011) for 7 days. To assess the proliferative and neurogenic response of ENS cells, animals were also exposed to EdU during the last 3 days of LY treatment (Fig. 7A). As expected, LY treatment of *Tg(her4.3:EGFP)* zebrafish resulted in rapid loss of GFP signal from the gut (Suppl. Fig. 8). Although this experiment confirmed that *Tg(her4.3:EGFP)* is a *bona fide* target of canonical Notch signalling in the ENS, it precluded the use of this transgene as a marker and lineage tracer of the EGC response to LY treatment. Therefore, we applied LY and EdU to *Tg(sox10:Cre;Cherry)* animals and analysed the entire population of non-neuronal ENS cells at the end of the LY/EdU treatment period (t0). As shown in Fig. 7B, Notch inhibition in 3-4 month old *Tg(sox10:Cre;Cherry)* zebrafish resulted in a dramatic increase in the percentage of non-neuronal ENS cells incorporating EdU (Cherry^+^HuC/D^-^EdU^+^) (control: 0.0387±0.21%; LY: 15.6±17.0%, p=2.67×10^-7^). A robust proliferative response of non-neuronal ENS cells was also observed in 6 month old *Tg(sox10:Cre;Cherry)* animals (control: 0.832±1.87%; LY: 6.95±8.2%, p=1.98×10^-5^) (Fig. 7D). Interestingly, LY treatment also resulted in increased enteric neurogenesis (Cherry^+^HuC/D^+^EdU^+^ cells) in both 3 month old (control: 0.0330±0.18%; LY: 2.12±7.8%, p=3.70×10^-4^) and 6 month old (control: 0.0652±0.22%; LY: 1.56±3.8%, 3.81 x 10^-4^) animals (Fig. 7C, E). Taken together, these experiments demonstrate that, similar to pallial RGCs (Alunni et al., 2013), the proliferation and neuronal differentiation of zebrafish EGCs are regulated by Notch signalling.

**Figure 7.**
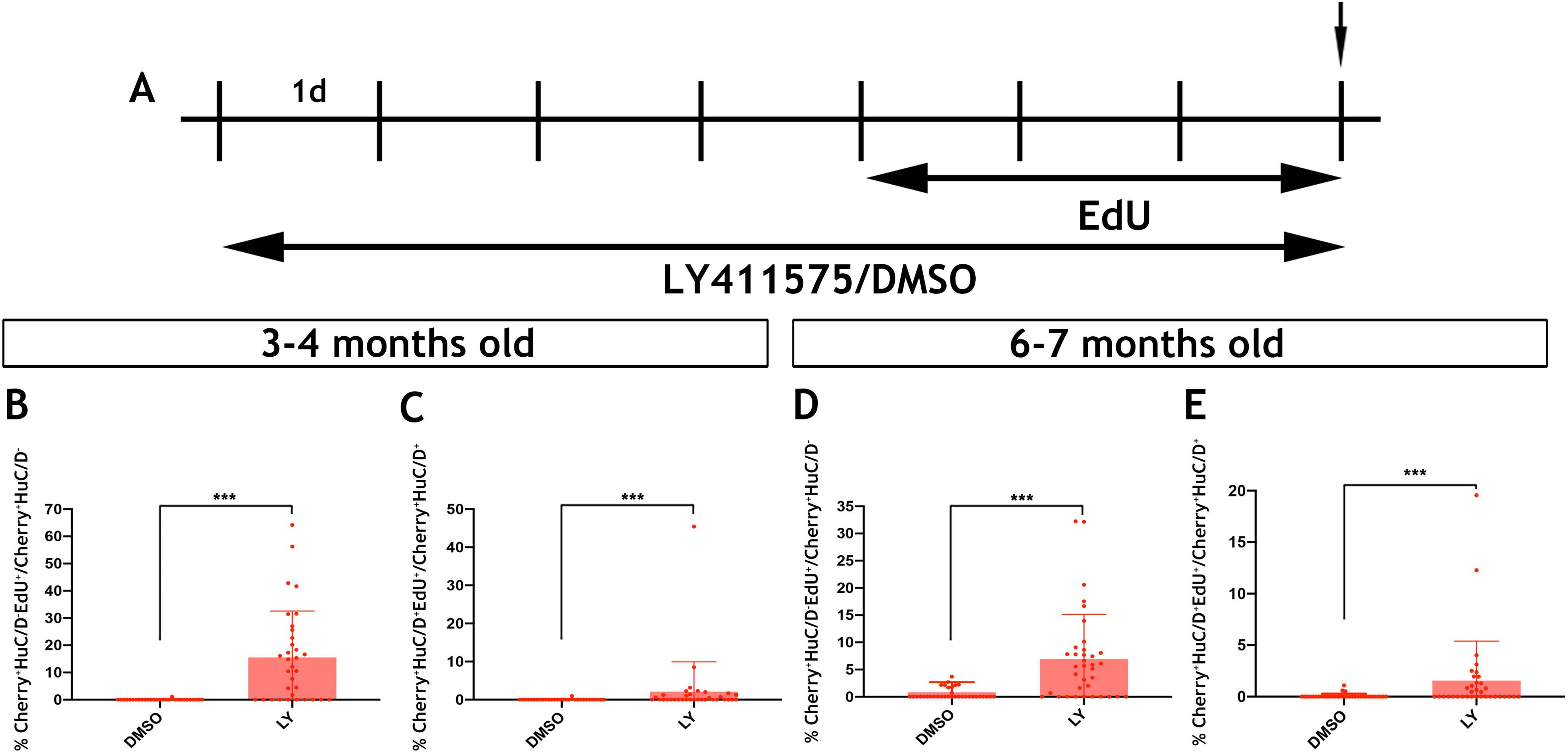
Notch signalling regulates the activation and differentiation of zebrafish EGCs. (**A**) Schematic representation of experimental protocol for LY/EdU treatment. (**B-E**) Quantification of the effect of Notch inhibition on the proliferation (B and D) and neuronal differentiation (C and E) of EGCs in 3-4 month old (B and C) and 6-7 month old (D and E) animals. Data are given as mean ±SD. *** P<0.001.

## Discussion

Here, we characterise the non-neuronal compartment of the zebrafish ENS and identify both familiar and unexpected properties of EGCs in teleosts. Specifically, we demonstrate that markers commonly used for the identification of peripheral glial cells in higher vertebrates are not detected in zebrafish EGCs, but that EGCs share morphological features and gene expression programmes with their mammalian counterparts. However, in contrast to mammalian enteric glia, but in accordance with the properties of brain RGCs, the population of zebrafish EGCs is dynamic, undergoing self-renewing proliferation and neuronal differentiation during homeostasis, which are regulated by Notch signalling. Our findings highlight the neural precursor potential of vertebrate enteric glia *in vivo* and reveal previously unanticipated similarities to brain NSCs.

Earlier histological studies demonstrated that mammalian enteric glia are remarkably similar to protoplasmic astrocytes and express the intermediate filament GFAP, a characteristic astrocytic marker (Jessen and Mirsky, 1980; Ruhl, 2005). Further EM analysis revealed diagnostic ultrastructural characteristics of intestinal neuroglia networks in rodents (Gabella, 1981). Extending these early reports, we and others have identified four morphological subtypes of mammalian enteric glia, which correlate with their position in the gut and relative to the ganglionic network in the gut wall (Boesmans et al., 2015; Gulbransen and Sharkey, 2012). Our current experiments demonstrate that all cardinal morphological and ultrastructural features ascribed to mammalian enteric glia are also found in the *Tg(her4.3:EGFP)^+^* non-neuronal compartment of the zebrafish ENS, thus providing strong evidence that it represents the EGC lineage of the teleost ENS. Our failure to detect glia markers commonly used to identify mammalian enteric glia (such as GFAP and S100b) indicates that the expression of these genes may not be integral to the genetic programmes operating in the vertebrate ENS, but rather signifies dynamic physiological states of EGCs adopted in response to specialised local cues. In support of this idea, GFAP is dynamic and is normally detected in a subpopulation of mammalian EGCs *in vivo* (Boesmans et al., 2015) and expression of GFAP and S100b is enhanced in primary cultures of human enteric glia challenged with pro-inflammatory stimuli (Cirillo et al., 2011). It would be interesting to determine whether these glia markers are also upregulated in zebrafish EGCs following inflammatory pathology, infection or injury.

Despite the failure to detect canonical glia marker expression, our transcriptomic analysis of zebrafish EGCs revealed a considerable overlap in the gene expression profile of teleost and mammalian enteric glia. Among the genes expressed by both lineages are those encoding the early NC cell markers *sox10*, *foxd3* and *plp1*, as well as the stem cell regulators *sox2* and *ptprz1a/b*. The roles of these genes have been studied extensively in the context of neural development (*sox10*, *foxd3*, *sox2*) and stem cell dynamics (*sox2*, *ptprz1a/b*), but their potential contribution to the homeostasis and function of enteric glia in adult animals remains unknown. We suggest that the shared gene expression modules we have identified between teleost and mammalian enteric glia represent evolutionary conserved regulatory programmes that are critical for ENS homeostasis and highlight the potential of vertebrate EGCs to serve as neurogenic precursors.

One of the unexpected findings of our work is the relatively small size of the non-neuronal compartment in the zebrafish ENS relative to its mammalian counterpart. A series of studies demonstrating that glial cells regulate synaptic activity of CNS neural circuits have led to the suggestion that the enhanced capacity of the higher vertebrate brain for neural processing has been fuelled during evolution by the increased number, size and complexity of astrocytes (Han et al., 2013; Oberheim et al., 2006). Perhaps the higher number of enteric glia in mammals relative to teleosts, may also reflect an increase in the functional complexity of intestinal neural circuits during vertebrate evolution and an enhanced scope of EGCs in the regulation of the complex gut tissue circuitry that maintains epithelial cell homeostasis, host defence and healthy microbiota (Grubisic and Gulbransen, 2016).

Several reports have documented that peripheral glial cells can acquire properties of neural crest stem cells (NCSCs) and give rise to diverse cell types. For example, Schwann cell precursors (SCPs) associated with growing nerves in mammalian embryos, in addition to generating the Schwann cell lineage of adult animals, also function as multipotent progenitors giving rise to diverse cell types, including mesenchymal and neuroendocrine cells, parasympathetic neurons and melanocytes (Parfejevs et al., 2018; Petersen and Adameyko, 2017). Echoing the developmental potential of SCPs, ENS progenitors already expressing molecular markers attributed to EGCs are also capable of generating enteric neurons and mature enteric glia (Cooper et al., 2016; Cooper et al., 2017; Lasrado et al., 2017). In addition to these studies, a growing body of evidence indicates that NCSC properties can be acquired by peripheral glia cell lineages from adult animals, including Schwann cells, glia of the carotid body and EGCs (Jessen et al., 2015; Pardal et al., 2007). However, it is generally thought that the reprogramming of differentiated glial cells into a NCSC-state is induced by injury, infection or other types of stress, including tissue dissociation and culture. Thus in mammals, EGCs can undergo limited neurogenesis in response to chemical injury to the myenteric plexus, pharmacological activation of serotonin signalling or bacterial gut infection (Belkind-Gerson et al., 2017; Joseph et al., 2011; Laranjeira et al., 2011; Liu et al., 2009). By providing evidence that zebrafish EGCs, in addition to their *bona fide* role as glial cells, also serve as constitutive ENS progenitors *in vivo*, our studies argue that the neurogenic potential of mammalian enteric glia disclosed under conditions of injury and stress, reflects an earlier evolutionary state of anamniote vertebrates, in which the same cell type exhibited properties of neural progenitors and mature glia. Although it is currently unclear whether neurogenic potential is a unique property of teleost EGCs, we speculate that peripheral glia lineages in lower vertebrates represent NCSCs that retain their developmental options but adjust to the cellular environment they reside in by acquiring additional specialised functions that contribute to local tissue function and homeostasis. Understanding the transcriptional and epigenetic mechanisms that underpin retention of the NCSC character and simultaneously allow novel functional adaptations during ontogenesis represents an exciting challenge of fundamental biology with practical implications. For example, identification of the molecular mechanisms that drive neuronal differentiation of enteric glia *in vivo*, will facilitate strategies to harness the intrinsic neurogenic potential of mammalian EGCs and restore congenital or acquired deficits of intestinal neural circuits.

By subsuming features of both neural progenitors and glial cells, zebrafish EGCs show remarkable and unexpected parallels to RGCs, NSCs that are distributed widely in teleost brain, reflecting its pronounced neurogenic and regenerative potential (Alunni and Bally-Cuif, 2016; Than-Trong and Bally-Cuif, 2015), and take on functions normally attributed to astrocytes (Lyons and Talbot, 2014). The parallels of RGCs and EGCs are likely to extend beyond a cursory parity imposed by the demands of the resident organs (brain and gut) for continuous growth and specialised glia function, and apply to specific cellular and molecular mechanisms controlling their homeostasis and differentiation. Thus, although the majority of RGCs and EGCs remain quiescent under physiological conditions ((Alunni et al., 2013) and this study), similar fractions enter the cell cycle and undergo neuronal differentiation to ensure the long-term maintenance of the original cell populations and the generation of new neurons to cater for the physical growth and plasticity of local neural networks (Supplementary Figure 9). Notch signalling and its target genes control the dynamics of NSCs in vertebrates (Chapouton et al., 2010; Imayoshi et al., 2010) and differential activity of Notch receptors regulates the proliferation and differentiation of RGCs in the germinal zones of the zebrafish brain (Alunni et al., 2013; Than-Trong et al., 2018). Notch signalling has also been implicated in the development of the mammalian ENS by inhibiting the intrinsic neurogenic programme of ENS progenitors (Okamura and Saga, 2008). The demonstration that the Notch activity reporter *Tg(her4.3:EGFP)* is activated in ENS progenitors shortly after they invade the gut and initiate neurogenic differentiation suggests a similar role of Notch signalling in the development of the zebrafish ENS, namely attenuation of the strong neurogenic bias of early ENS progenitors acquired as they approach and enter the foregut and indicated by induction of strong neurogenic transcription factors, such as Phox2B and Ascl1 (Charrier and Pilon, 2017). Although the relevant Notch signalling components remain to be identified, our findings argue that activation and differentiation of EGCs in adults is also under the control of Notch signalling pointing to further fundamental similarities in the mechanisms controlling the homeostasis of CNS and ENS in vertebrates. The hierarchical relationships of cell types in the non-neuronal compartment of the zebrafish ENS and the potential regional differences in the dynamics of EGCs in zebrafish gut remain to be characterised. Nevertheless, the systematic comparison of the molecular properties of mammalian and teleost enteric glia are likely to identify cell-intrinsic molecular cascades responsible for the differences in the *in vivo* neurogenic potential of the two lineages, thus offering opportunities for the repair of defective gastrointestinal neural networks in mammals by activating endogenous EGCs.

## Materials and Methods

### Animals

All animal experiments were carried out in compliance with the Animals (Scientific Procedures) Act 1986 (UK) and in accordance with the regulatory standards of the UK Home Office (Project Licence PCBBB9ABB). Experimental protocols were approved by the local Animal Welfare and Ethical Review Body (AWERB) of the Francis Crick Institute. Zebrafish stocks were maintained as described (Heanue et al., 2016a; Westerfield, 2000). Embryos and larvae were maintained and staged as described (Heanue et al., 2016a), while embryos used for time lapse were reared in 0.2mM PTU from 24hpf to inhibit melanisation, as described (Westerfield, 2000). Transgenic and mutant lines used were as follows: *Tg(SAGFF234A)* (Asakawa et al., 2008; Kawakami et al., 2010); *Tg(UAS:GFP)* (Kawakami et al., 2010), *Tg(−4.7sox10:Cre)* (Rodrigues et al., 2012), *Tg(*β*actin-LoxP-STOP-LoxP-hmgb1-mCherry)* (Wang et al., 2011), *ret^hu2846^* (Knight et al., 2011), *Tg(gfap:GFP)* (Bernardos and Raymond, 2006), *Tg* (−*3.9nestin:GFP*) (Lam et al., 2009), *Tg(her4.3:EGFP)* (Yeo et al., 2007), *Tg(SAGFF217B)* (Kawakami et al., 2010). Note that the *Tg(her4.3:EGFP)* designation is the current ZFIN reference for this transgene, however it is also variously referred to as *Tg(her4:EGFP)* (Yeo et al., 2007) or *Tg(her4.1GFP)* (Kizil et al., 2012). *her4.3* is one of 6 (of 9) mammalian orthologues of mammalian *Hes5* found in tandem duplication on chromosome 23 of the zebrafish genome (Zhou et al., 2012). The stable *Tg(UAS:mmCherry)* line was generated by Tol2 transgenesis: co-microinjection of TOL2 transposase with a construct containing membrane-mCherry (mmCherry) downstream of two copies of the Gal4 recognition sequence UAS, with bicistronic α crystalinP:RFP cassette enabling red eye selection of carriers, as described previously (Gerety et al., 2013). Genotyping was done based on the lines’ previously described distinct fluorescent patterns, or by PCR in the case of *Tg(ret ^hu2846/+^)*, as described (Knight et al., 2011).

### Immunohistochemistry

Immunohistochemistry was performed as previously described on whole-mount embryos/larvae or whole-mount adult intestines and brains (Heanue 2016). Primary antibodies used were as follows: HuC/D (mouse, ThermoFisher A21272, 1:200), Cherry (goat, Antibodies online ABIN1440057, 1:500), GFP (chick, Abcam ab13970, 1:500), S100β (rabbit, Dako Z0311, 1:500), mGFAP (rabbit, Sigma G9269, 1:500), zGFAP (rabbit, Genetex GTX128741, 1:500), zrf-1 (mouse, Abcam ab154474, 1:200), BFABP (Merck ABN14, 1:500), AcTu (mouse, Sigma T6793, 1:1000), MCM5 antibody (1:500, kindly provided by Soojin Ryu, Max Planck Institute for Medical Research, Heidelberg, Germany) and appropriate secondary antibodies conjugated to AlexaFluor 405, 488, 568 and 647 were used for visualisation (Molecular Probes). EdU was developed using the EdU Click-it kit following the manufacturer’s instructions and combined with fluorophores Alexa555 or Alexa647 (C10337 and C10339). For MCM5 labelling, antigen retrieval was required to expose the epitope. Briefly, after immunostaining for GFP, antigen retrieval with Citrate buffer (pH6.0) was performed. All tissues were mounted on Superfrost Plus™ slides with Vectashield Mounting Media with/without DAPI (H1200/H1000, respectively). Immunohistochemistry images were captured on a Leica CM6000 confocal microscope or an Olympus FV3000 confocal microscope, with standard excitation and emission filters for visualising DAPI, Alexa Flour 405, Alexa Flour 488, Alexa Flour 568 and Alexa Flour 647. Cell counting analysis was carried out using the Cell Counter plugin on Fiji and images processed with Adobe Photoshop 8.

### Purification of ENS nuclei from adult gut muscularis externa

Adult *Tg(sox10:Cre;Cherry)* zebrafish intestines were first dissected, then cut along their length and immersed in HBSS (no calcium, no magnesium, (ThermoFisher 14170088) containing 20mM EDTA and 1% Penicillin/Streptomycin (ThermoFisher, 15140122) for 20-25 minutes at 37°C until the epithelia cell layer was seen to begin detaching from the overlying muscularis externa, evident by clouding of the HBSS solution. After several washes in PBS (ThermoFisher 14190094), the tissue was placed under a dissecting microscope and the muscularis externa was grasped in forceps and agitated briefly to detach any remaining associated epithelial cells. Muscularis externa was tranfered to a fresh tube and purification of nuclei was performed essentially as described (Obata et al., 2020). Briefly, dounce homogenization was performed in lysis buffer (250mM sucrose, 25mM KCl, 5mM MgCl_2_, 10mM Tris buffer with pH8.0, 0.1mM DTT) containing 0.1% Triton-X, cOmplete™ EDTA-free protease inhibitor (Sigma-Aldrich) and DAPI. The homogenate was filtered to remove debris and centrifuged to obtain a pellet containing the muscularis externa nuclei. For flow cytometric analysis, doublet discrimination gating was applied to exclude aggregated nuclei, and intact nuclei were determined by subsequent gating on the area and height of DAPI intensity. Both mCherry^+^ and mCherry^-^ nuclear populations (termed Cherry^+^ and Cherry^-^ in text and figures) were collected directly into 1.5mL tube containing Trizol LS reagent (Invitrogen) using the Aria Fusion cell sorter (BD Biosciences). The obtained FCS data were further analysed using FlowJo software version 10.6.1. For each replicate, sorted cells from an average of 30 adult guts were pooled, containing approximately 30,000 mCherry^+^ or mCherry^-^ nuclear populations.

### RNA-sequencing and Bioinformatic analysis

RNA was isolated from nuclei populations using the PureLink RNA Micro Kit (Invitrogen #12183016), according to the manufacturer’s instructions. Double stranded full-length cDNA was generated using the Ovation RNA-Seq System V2 (NuGen Technologies, Inc.). cDNA was quantified on a Qubit 3.0 fluorometer (Thermo Fisher Scientific, Inc.), and then fragmented to 200bp by acoustic shearing using Covaris E220 instrument (Covaris, Inc.) at standard settings. The fragmented cDNA was then normalized to 100ng, which was used for sequencing library preparation using the Ovation Ultralow System V2 1-96 protocol (NuGen Technologies, Inc.). A total of 8 PCR cycles were used for library amplification. The quality and quantity of the final libraries were assessed with TapeStation D1000 Assay (Agilent Technologies, Inc.). The libraries were then normalized to 4 nM, pooled and loaded onto a HiSeq4000 (Illumina, Inc.) to generate 100 bp paired-end reads.

### Bioinformatics Method Summary RNA-Sequencing-analysis

Sequencing was performed on an Illumina HiSeq 4000 machine. The ‘Trim Galore!’ utility version 0.4.2 was used to remove sequencing adaptors and to quality trim individual reads with the q-parameter set to 20 (https://www.bioinformatics.babraham.ac.uk/projects/trim_galore/). Then sequencing reads were aligned to the zebrafish genome and transcriptome (Ensembl GRCz10 release-89) using RSEM version 1.3.0 (Li and Dewey, 2011) in conjunction with the STAR aligner version 2.5.2 (Dobin et al., 2013). Sequencing quality of individual samples was assessed using FASTQC version 0.11.5 (https://www.bioinformatics.babraham.ac.uk/projects/fastqc/) and RNA-SeQC version 1.1.8 (DeLuca et al., 2012). Differential gene expression was determined using the R-bioconductor package DESeq2 version 1.14.1 (Love et al., 2014)(R Development Core Team (2008). R: A language and environment for statistical computing. R Foundation for Statistical Computing, Vienna, Austria. ISBN 3-900051-07-0, URL http://www.R-project.org). Gene set enrichment analysis (GSEA) was conducted as described in (Subramanian et al., 2005). For conversion from mouse to zebrafish gene names we used NCBI homologene (ftp://ftp.ncbi.nih.gov/pub/HomoloGene/current/homologene.data), with a manually curated addition as shown in Supplementary Table 3.

### Fluorescence in situ hybridization

Adult zebrafish intestines from *Tg(sox10:Cre;Cherry*) or *Tg(her4.3:EGFP)* were first dissected, then cut along their length, pinned to a silguard plate and immersed in HBSS (ThermoFisher 14170088) containing 20mM EDTA and 1% penicillin/streptomycin (ThermoFisher, 15140122) for 20-25 minutes at room temperature to detach the epithelia layer. After several washes in PBS (ThermoFisher 14190094), the epithelia was manually teased away with forceps. After washing in PBS, 4% PFA was added to the plate with pinned tissue to fix overnight at 4°C. Fluorescence in situ hybridization was then performed using the Advanced Cell Diagnostics RNAscope® Fluorescent Multiplex Kit (ACD #320850), according to manufacturer’s specification and essentially as described (Obata et al., 2020). Briefly, tissue was washed in PBS, dehydrated through an ethanol series and then incubated with RNAscope® Protease III for 25 minutes. Tissue was incubated overnight at 40°C in a HybeOven with customized probes (*sox10*, *foxd3*, *ret*). The next day, the tissue was washed twice with Wash Buffer before hybridization the with pre-amplifier, the appropriate amplifier DNA (Amp 1-FL, Amp 2-FL and Amp 3-FL) and appropriate fluorophores (Amp4 Alt A-FL/AltC-FL) at 40°C for 15-30 minutes, as per the manufacturer’s instructions. Tissues were then processed for immunohistochemistry and mounted directly onto Superfrost Plus™ slides (ThermoFisher Scientific #10149870) Vectashield Mounting Media without DAPI (VectorLabs H1000). Image were captured on a Leica CM6000 confocal microscope or an Olympus FV3000 confocal microscope, with standard excitation and emission filters for visualising DAPI, Alexa Flour 405, Alexa Flour 488, Alexa Flour 568 and Alexa Flour 647 and images processed with Adobe Photoshop 8.

### Correlative Light and Electron Microscopy

Intestines were dissected from *Tg(her4.3:EGFP;SAGFF217B;UAS:mmCherry)* adult animals and fixed in 4% formaldehyde 0.1% glutaraldehyde in phosphate buffer (PB) overnight at 4°C. Subsequently, the intestines were sectioned to 150µm on a Leica vibratome, and stored in 2% formaldehyde in PB. Super-resolution images of mid-gut sections were mounted in PB on SuperFrost Plus™ slides and imaged with an inverted Zeiss 880 confocal microscope with AiryScan, using standard emission and excitation filters for EGFP and mmCherry. A low magnification overview image was acquired using a 20× objective before 2-3 regions of interest (ROI) were identified per section that contained at least one EGFP^+^ cell of interest.

The Airyscan was aligned for EGFP and mmCherry using an area outside of the ROIs where both fluorophores were identified. After Airyscan alignment, the ROIs were captured using a x63 glycerol objective and pixel size, z-depth and zoom (>1.8x) were defined by Nyquist’s theorem. Once fluorescence microscopy was completed, the vibratome slices were further fixed in 2.5% glutaraldehyde and 4% formaldehyde in 0.1 M phosphate buffer (pH 7.4), and processed according to the method of the National Centre for Microscopy and Imaging Research (Deerinck TJ, Bushong EA, Thor A, Ellisman MH (2010) NCMIR methods for 3D EM: a new protocol for preparation of biological specimens for serial block face scanning electron microscopy https://ncmir.ucsd.edu/sbem-protocol) before flat embedding between sheets of Aclar plastic.

### SBF SEM data collection and image processing

Serial blockface scanning electron microscopy (SBF SEM) data was collected using a 3View2XP (Gatan, Pleasanton, CA) attached to a Sigma VP SEM (Zeiss, Cambridge). Flat embed vibratome slices were cut out and mounted on pins using conductive epoxy resin (Circuitworks CW2400). Each slice was trimmed using a glass knife to the smallest dimension in X and Y, and the surface polished to reveal the tissue before coating with a 2 nm layer of platinum. Backscattered electron images were acquired using the 3VBSED detector at 8,192*8,192 pixels with a dwell time of 6 µs (10 nm reported pixel size, horizontal frame width of 81.685 µm) and 50 nm slice thickness. The SEM was operated at a chamber pressure of 5 pascals, with high current mode inactive. The 30 µm aperture was used, with an accelerating voltage of 2.5 kV. A total of 1,296 images were collected, representing a depth of 64.8 µm, and volume of 432,374 µm^3^. Downstream image processing was carried out using Fiji (Schindelin et al., 2012). The images were first batch converted to 8-bit tiff format, then denoised using Gaussian blur (0.75 pixel radius), and resharpened using two passes of unsharp mask (10 pixel radius 0.2 strength, 2 pixel radius 0.4 strength), tailored to suit the resolution and image characteristics of the dataset. Image registration was carried out using the ‘align virtual stack slices’ plugin, with a translation model used for feature extraction and registration. The aligned image stacks were calibrated for pixel dimensions, and cropped to individual regions of interest as required. To generate a composite of the two volumes, Bigwarp (Bogovic JA, 2015; Russell et al., 2017) was used to map the fluorescence microscopy volume into the electron microscopy volume which was reduced in resolution to isotropic 50 nm voxels to reduce computational load. The multi-layered cellular composition of the tissue was noted to have caused substantial non-linear deformation during processing of the sample for electron microscopy when compared to prior fluorescence microscopy. After exporting the transformed light microscopy volume from Bigwarp, a 2 pixel Gaussian blur was applied, the datasets were combined, and the brightness/contrast adjusted for on-screen presentation. False coloured images were composed by annotating separate semi-transparent layers in Adobe Photoshop CC 2015.5 with reference to prior fluorescence microscopy and 3-dimensional context within the image stack. Only processes that could be clearly tracked through the volume from definitively marked cell bodies were coloured.

### Time lapse imaging of zebrafish larvae

Embryos were raised in 0.2mM PTU, lightly anaesthetised with 0.15mg/ml Tricaine, and mounted into embryo arrays and overlayed with 0.6% low melt temperature agarose in embryo media essentially as described (Heanue et al., 2016a; Megason, 2009). Once set, the mould was overlaid with embryo media containing 0.15mg/ml Tricane and 0.15M PTU, and was replaced at least every 24hours. Larvae were imaged using a Leica CM6000 confocal microscope, with a 20X water dipping objective. Standard excitation and emission filters were used to visualise EGFP and mmCherry expression. For each individual embryo, 33 z-stacks (z thickness 2.014 µm) were collected at a frame rate of 602s, for 40.333 hours. Cells from the time-lapse recordings were tracked manually using the MTrack2 plugin on Fiji. To correct for growth or movement during the imaging process a reference point was taken, for each animal, as the point the anterior most spinal nerve, visible in the field of view, touched the gut. All calculated distances were given relative to this reference point.

### EdU labelling

To label proliferative cells, adult zebrafish were kept in system water with 1mM of EdU (0.05% DMSO) for 72 hours at a density of 4 zebrafish/litre. During chase periods adult zebrafish were kept in system water, which was changed every 2-3 days.

### Mathematical modelling

Since the zebrafish ENS is largely confined to the myenteric plexus, and hence the zebrafish ENS resides within a two dimensional plane, therefore, only X and Y coordinates were used for subsequent analysis. Each image covered a 450 μm-450 μm area and XY coordinates of individual cells were taken as the centre of the nucleus and obtained from the CellCounter plugin for Fiji. We first estimated the density of specifically labelled cells at several distances around every cell type of interest using confocal images with an area 45μ density was estimated in circles of increasing radius, r ∈ (20, 30, 40, .., 100, 150, ..,500 m), by dividing the number of cells within the circle by the surface area of the circle included within the image. When the radius was larger than the distance of the cell to the image edge, the area of the circle section overlapping with the image was numerically estimated by Monte Carlo simulation methods. We performed 50 Monte Carlo simulations for each confocal image with the observed number of cells of each phenotype in rearranged locations, according to a uniform distribution, on the 450 μm x 450 μm square area. Cell densities were estimated for each simulation as described above. To compare the recorded and simulated densities, we estimated the 90% confidence interval for simulated cell density under the assumption of cell homogeneity by fitting the gamma distribution function to the simulated values. When the average of the measured cell densities lied outside the 90% confidence interval, the observed spatial location was considered to be a non-chance event in a homogenous mixture of cells.

### Notch inhibition

Notch signalling was inhibited by immersion with 10µM LY411575 (Cambridge bioscience, 16162) (0.04% DMSO) in the system water, and was changed every 2-3 days, control zebrafish were incubated with the equivalent concentration of DMSO (0.04%).

### Statistics

Statistical analyses was performed using R 3.3.1. Due to the non-normality of most of the data, all comparisons were carried out using a two-sided Mann-Whitney non-parametric test. Resultant p-values were corrected for multiple testing using the Benjamini-Hochberg method as implemented by the p.adjust() function. A Pvalue of ≤ 0.05 was deemed to be significant and in figures designation of graded significance was as follows: P>0.05 (ns = non-significant), P ≤ 0.05 (*), P ≤ 0.01 (**), P ≤ 0.01 (***).

## Acknowledgments

We are grateful to Laure Bally-Cuif, Alessandro Alluni, Emmanuel Than-Trong for providing *Tg(her4.3:EGFP)* transgenic fish and specialist knowledge, to Donald Bell for expert advice in time-lapse imaging experiments, to the Aquatics BRF staff for fish husbandry, to the Flow Cytometry STP for FACS cell sorting, to the Advanced Sequencing Facility STP for library prep and sequencing, and to Joe Brock for scientific illustrations.

## Supplementary Figure Legends

**Supplementary Figure 1. ENS lineage tracing shows that there is a small non-neuronal lineage that is not detectable using antibodies for the canonical glial markers BFABP, GFAP nor with transgenic reporters.** (A) Using the *Tg(SAGFF234A;UAS:GFP)* line at 7dpf to label the ENS lineage with GFP (green), we observe that the majority of these cells are HuC/D^+^ neurons (cyan). (B) High magnification view of box in A, with arrows denoting the GFP^+^HuC/D^+^ ENS neurons and arrowheads indicating GFP^+^HuC/D^-^ non-neuronal ENS cells. (C) Quantification of neuronal (blue) and non-neuronal (green) populations within the 7dpf ENS lineage reveals that the majority of cells are neurons (93.7% ±3.0 vs 6.3% ±3.0). (D-I). The larval zebrafish ENS is not labelled with BFABP and GFAP antibodies. (D) BFABP (green) fails to mark EGCs in the 7dpf intestine despite HuC/D neurons (red) being readily detected. (E) The mammalian GFAP antibody (mGFAP, green) does not detect cells in the 7dpf gut, despite HuC/D positive neurons being detectable (red). Instead, mGFAP fibres are seen descending toward, but not entering, the gut (arrowheads). (F-G) An antibody raised against zebrafish GFAP (zGFAP) detects abundant circumferential fibres in the 7dpf gut (red, arrows), positioned near HuC/D^+^ ENS neurons (blue). However identical staining is observed in wild type larvae that contain ENS neurons (F) and *ret^hu2846/hu2846^* which lack an ENS due to a mutation in the Ret receptor tyrosine kinase and a failure of ENS progenitors to colonise the gut (G) (HuC/D^+^ neurons only present in F, blue). (H-I) Immunostaining of 7dpf *Tg(SAGFF234A;UAS:GFP)* larvae with another GFAP antibody raised against zebrafish GFAP (zrf-1) also reveals abundant circumferential fibres (red, arrows), in a pattern indistinguishable between wild type larvae containing ENS neurons (green) (H) and *ret^hu2846/hu2846^* larvae lacking ENS neurons (green, I), indicating that these fibres are not associated with the ENS lineage. (J-N) Antibodies tested in the above experiments to detect ENS glial cells are able to successfully label CNS glial cells in the 7dpf spinal cord: S100b (J), BFABP (K), mGFAP (L), zrf-1 (M), zGFAP (N). (O) The expected pattern of GFP^+^ cells are detected within the spinal cord of 7dpf *Tg(gfap:GFP)* larvae. (P-R) Analysis of adult gut tissue using a variety of antibody and transgenic tools used to identify CNS glial cells. (P) BFABP is not detected in the adult gut despite HuC/D (red) identifying HuC/D^+^ enteric neurons. (Q) Although signal is detected in the adult gut using the zGFAP antibody (green), the striated signal is not found in cell bodies, nor is it clearly associated with HuC/D neurons (red) and the staining pattern is reminiscent of the non-ENS associated staining seen at 7dpf. (R) GFP^+^ cells are not observed in adult *Tg(−3.6nestin:GFP)* gut tissue, despite the ready detection of HuC/D^+^ neurons (red). 50µm scale bars in merge panels (A, D-I, P-R) or single colour images (J-O).

**Supplementary Figure 2. Transcriptional profiling of adult zebrafish ENS nuclei identifies profiles indicative of both neurons and glia** (A) A representative FACS plot showing nuclei from the muscularis externa of adult *Tg(sox10:Cre;Cherry)* zebrafish guts gated on single intact DAPI^+^ nuclei. mCherry^+^ nuclei were collected, representing less than 1% of the starting population. An equivalent number of mCherry^-^ nuclei were also collected. (B) Principal component analysis of the adult gut transcriptomes reveals segregation of the samples by Cherry^+^ vs. Cherry^-^ expression (30% of variability explained in PC1, 13% in PC2). (C-H) Analysis of the adult gut Cherry^+^ vs Cherry^-^ transcriptomic data by comparison to previously published data and publicly available reference data. The adult gut Cherry^+^ vs Cherry^-^ transcriptomic data was filtered to select those genes with log fold-change > 0 (in Cherry^+^ vs Cherry^-^) and with p-value < 0.05. The resulting set is enriched for statistically significant zebrafish ENS-associated genes. Cross-species comparisons (zebrafish to mouse) utilise publicly available homology assigning resources (see methods). (C) Gene set enrichment analysis shows that GO Biological Processes enriched in the Cherry^+^ population include nervous system associated terms. (D-E) Enrichment plots of representative gene sets (D) Synaptic Signalling and (E) Neuron cell-cell adhesion shows enrichment in Cherry^+^ samples. (F) Clustered heat map showing expression of a list of genes enriched in zebrafish larval ENS neurons (from Roy-Carson, 2017) that is analysed in our adult zebrafish gut transcriptomic data. We observe that >750 of these neural expressed genes are enriched in the Cherry^+^ samples relative to Cherry^-^ samples (Supplementary Table 1), and these are candidate adult ENS neuron-associated genes. These include *phox2bb*, *phox2a*, *ret*, *elavl3*, *elavl4*, *vip*, and *nmu*. (G) Clustered heat map showing the top 25 genes identified as enriched in mammalian Plp1^+^ glial cells (Rao, 2015) that have zebrafish orthologues and which are upregulated in Cherry^+^ vs Cherry^-^ samples, revealing 9 candidate zebrafish ENS glial cell-associated genes. (H) Clustered heat map showing expression of genes in the adult zebrafish ENS transcriptome after removing genes associated with zebrafish ENS neurons (from C, above). Over 600 unique genes are identified (Supplementary Table 2), which are candidate adult ENS non-neuronal or ENS glial cell-associated genes. These include *sox10*, *foxd3*, *sox2*, *plp1b*, *ptprz1a* and *ptprz1b*.

**Supplementary Figure 3. Her4.3GFP transgenic line identifies cells with morphologies indicative of distinct** subtypes of EGCs Immunohistochemistry of adult guts from of *Tg(her4.3:EGFP)* allow characterization the cellular morphology of GFP^+^ cells and comparisons to mammalian EGC subtypes (Boesmans et al., 2015). (A) GFP expressing cells (green) show close association with neurons, which express HuC/D in cell bodies and AcTu in cell processes (red). Inset shows high magnification view of boxed region, marking neurons (asterisks), GFP expression (arrowhead), and highly branched GFP-expressing cellular processes (arrows). (B-E) Four distinct morphological cell types can be observed in *Tg(her4.3:EGFP)*^+^ cells: (B) GFP^+^ cells in the myenteric layer (arrowhead) with processes that appear to wrap around HuC/D^+^ cell bodies (red, asterisk), similar to Type 1 mammalian EGCs (inset), (C) GFP^+^ cells in the myenteric layer (arrowhead) with elongated processes (arrow) that follow AcTu^+^ neuronal processes (red), similar to Type 2 mammalian EGCs (inset), (D) GFP^+^ cells close to the mucosal layers (arrowheads), such as mucosal epithelia (ep, with DAPI highlighted nuclei in grey), similar to mammalian Type 3 EGCs (inset), and (E) Bipolar GFP^+^ cells within the muscle layers (arrowhead), associated with AcTu^+^ neuronal fibres (red, arrow), similar to Type 4 mammalian EGCs (inset). Inset pictures adapted from (Boesmans et al., 2015). Scale bars in merge panels: 50µm (A) and 10µm (B-E).

**Supplementary Figure 4. Correlative light-electron microscopy identifies glial like features of *Tg(her4.3:EGFP* expressing cells** (A) A subpopulation of HuC/D^+^ ENS neurons (green) are highlighted by *Tg(SAGFF217;UAS:mmCherry)*, and Cherry expression (red, arrows) fills both the cell bodies and the abundant processes of expressing cells (red). The remaining proportion on HuC/D^+^ cells (green) do not express Cherry (arrowhead). (B) Electron microscopy image of a section from a *Tg(her4.3:EGFP;SAGFF217;UAS:mmCherry)* gut with tissue layers denoted, false coloured to depict the position of the GFP^+^ cell shown in the super resolution image shown in inset. Note the neuron and axons in this section are not Cherry^+^ neurons. (C) High magnification view of the boxed region, showing crenelated nuclei (arrows) and radial extensions that separate axon bundles (yellow arrowheads, asterisk denotes axon bundle), and many which contact the neuronal cell body (neuronal cell body denoted with N). Scale bars: 10µm (A,B) and 1µm (C).

**Supplementary Figure 5. Lineage analysis reveals that *Tg(her4.3:EGFP)* expressing cells are derived from the NC cell population that gives rise to the ENS** (A-C) Analysis of *Tg(her4.3:EGFP;SAGFF234A*;*UAS:mmCherry)* allows *her4.3:EGFP^+^* cells to be examined relative to the Cherry^+^ migrating NC cell population that colonises the gut. (A) At 54hpf, no GFP^+^ cells (green) are present in the gut and none are detected within the population of migrating NC cells (red), although NC cell-derived HuC/D^+^ ENS neurons are present at this time (blue). Single channels shown in high magnification view of boxed region. (B) At 72hpf, small numbers of weakly GFP-expressing cells (green, arrows) can be seen within the streams of NC cells colonising the gut (red). GFP^+^ cells are seen in proximity to HuC/D^+^ cells (blue). Note strongly GFP-expressing cells can be detected, but these cells do not form part of the NC cell migratory streams (red) and are outside of the gut (grey arrowhead), and are likely to be melanocytes. Single channels shown in high magnification view of boxed region. (C) At 4dpf, an increased number of both strongly and weakly GFP expressing cells (green, arrows) are found within the stream of migratory NC cells (red). Single channels shown in high magnification view of boxed region. (D) At 7dpf *Tg(her4.3:EGFP)* larvae GFP-expressing cells (green, arrowheads) are closely associated with, but distinct from, HuC/D^+^ positive neurons (red). Occasionally HuC/D is seen to overlap with cells expressing low levels of GFP (open arrowheads). Scale bars in merge panels: 50µm.

**Supplementary Figure 6. The *Tg(her4.3:EGFP)* cells are actively proliferating in adult homeostasis.** (A) *Tg(her4.3:EGFP)* zebrafish flattened intestines immunostained for GFP (green) and the cell-cycle marker MCM5 (red). Actively proliferating GFP^+^MCM5^+^ cells were observed (arrows) throughout the intestine. The majority of the GFP^+^ population remains quiescent (arrowheads). (B) Quantification of the percentage of GFP^+^MCM5^+^ cells over the total GFP^+^ population. Data are given as mean ± SD. Scale bar: 10µm in merge panel.

**Supplementary Figure 7. *Tg(her4.3:EGFP)* cells take up EdU and appear in doublets.** (A) Schematic of experimental design: Immersion of 3 month old adult *Tg(her4.3:EGFP)* zebrafish in 1mM EdU pulse for three days was followed by a return to normal zebrafish water. Animals were then culled after chase periods of 0 days (t0), 4 days (t4) or 11 days (t11) and analysed for EdU incorporation. (B-D) At 0 days chase, the majority of EdU labelled GFP^+^ (yellow) cells are found in doublets (two labelled cells in close proximity). These cells are either: (B) both expressing high levels of GFP (green, arrows), (C) appear with one high GFP expressing cell (arrow) and one low GFP expressing cell (arrowhead), (D) in larger groupings, where EdU labelling is associated with cells exhibiting lower levels of GFP expression (arrowhead) and not observed in high GFP expressing cells (arrows). Scale bars in merge panels: 10µm (B-D).

**Supplementary Figure 8. Notch inhibition leads to loss of GFP expression from the *Tg(her4.3:EGFP)* transgene.** (A) After 7 days of DMSO treatment the *Tg(her4.3:EGFP)* transgene (green) is clearly visible within the adult ENS, along with HuC/D^+^ neurons (red). (B) After 7days of treatment with the γ-secretase inhibitor LY411575 led to a specific reduction of *Tg(her4.3:EGFP)* expression was observed. Scale bars in merge panels: 50µm.

**Supplementary Figure 9: Working model of enteric glia acting as a source of neural progenitors in adult zebrafish during homeostatic conditions.** Given the similarities between *Tg(her4.3:EGFP)*^+^ EGCs and *Tg(her4.3:EGFP)*^+^ RGCs, we propose that like RGCs, EGCs may exist in two forms: *Tg(her4.3:EGFP)*^+^ quiescent EGCs (qEGCs) and *Tg(her4.3:EGFP)*^+^ activated EGC (aEGCs), the latter of which are proliferative and can take up EdU in our experiments (indicated in blue). We suggest that aEGCs are a self-renewing population, which may also revert to the quiescent state. The proliferative aEGC population can give rise to enteric neuronal progenitors (eNP; cells committed to the neurogenic lineage), which can retain EdU but are *Tg(her4.3:EGFP)*^-^ and will not yet express HuC/D. These cells would correspond to the Cherry^+^GFP^-^HuC/D^-^EdU^+^ cells quantified in Figure 6I, which increase during the EdU labelling period of our experiments. Finally, neural progenitors undergo full neuronal differentiation (eN), can be detected with HuC/D and are also EdU^+^ in our experiments. These cells correspond to the Cherry^+^GFP^-^HuC/D^+^EdU^+^ quantified in Figure 6H, which also increase during the course of our EdU labelling experiments.

**Supplementary Table 1: Table containing the order of heatmap genes and values for Supplementary** Figure 2F. Genes displayed in the heat map depicting the nRNASeq data of this study were selected as follows: genes with a logFC (Cherry^+^ vs Cherry^-^) > 0, padj (Cherry^+^ vs Cherry^-^) < 0.05 and an average TPM of 3. We intersected this selection with the 2561 genes identified in “Additional File 2: TableS1” of Roy-Carson et al., 2017 as upregulated in 7dpf phox2b:EGFP^+^ gut cells relative to EGFP^-^ gut. This selection highlights 758 genes. Gene names and Ensembl gene IDs found in column K.

**Supplementary Table 2: Table containing the order of heatmap genes and values for Supplementary** Figure 2H. Genes displayed in the heat map depicting the nRNASeq data of this study were selected as follows: genes with a logFC (Cherry^+^ vs Cherry^-^) > 0, padj (Cherry^+^ vs Cherry^-^) < 0.05 and an average TPM of 3. We removed from this list the genes found in Supplementary Table 1. This selection highlights 660 genes. Gene names and Ensembl gene IDs found in column K.

**Supplementary Table 3: Zebrafish orthologues of the mouse genes identified in Table 1** of Rao et al., 2015 PMID: 26119414 “Top 25 genes enriched in PLP1+ enteric glia”, generated using the ZFIN and Ensembl databases. Column A shows the zebrafish gene names of the orthologues of the mouse genes shown in Column B

**Supplementary Movie 1: Correlative light and electron microscopy** (CLEM) analysis of the adult *Tg(her4.3:EGFP;SAGFF234A;UASmmCherry)* gut Mapping of the super-resolution light microscopy volume into the cropped SBF SEM volume using Bigwarp confirmed the identification and localisation of EGFP+ non-neuronal cells and mmCherry+ neurons within a 3D region of interest from the midgut of *Tg(her4.3:EGFP;SAGFF217;UAS:mmCherry)* zebrafish. The EGFP+ cells and mmCherry+ neurons that were false coloured in figure 4 and supplementary figure 4 are indicated with green and red arrows, respectively, showing that each forms numerous complex extensions through the volume. Data is shown at 10 frames per second, with 100 nm pixels in XY (cropped to represent a horizontal frame width of 80.5 um) and 50 nm pixels in Z (representing a depth of 64.8 um).

**Supplementary Movie 2: Representative time-lapse image from a** *Tg(her4.3:EGFP;SAGFF234A;UASmmCherry)* embryo. Time-lapse imaging revealed that *Tg(her4.3:EGFP)^+^* cells (grey, white arrowheads) are found within the mmCherry^+^ neural crest cells (red) that are colonising the developing gut, but the EGFP^+^ cells appear behind the wavefront of migration (red arrowheads). Time given is shown as hh:mm from the start of recording. See methods for details.

**Supplementary Movie 3: Representative movie of *de novo* EGFP** expression in time-lapse movies from *Tg(her4.3:EGFP;SAGFF234A;UASmmCherry)* embryos. De-novo *her4.3:EGFP* transgene expression (grey) within the enteric nervous system (red) is observed during time lapse recordings of developing *Tg(her4.1:EGFP;SAGFF234A;UAS:mCherry)* embryos (arrow).

